# Exposure to *mycobacterium* remodels alveolar macrophages and the early innate response to *Mycobacterium tuberculosis* infection

**DOI:** 10.1101/2022.09.19.507309

**Authors:** Dat Mai, Ana Jahn, Tara Murray, Michael Morikubo, Pamelia N. Lim, Maritza M. Cervantes, Linh K. Pham, Johannes Nemeth, Kevin Urdahl, Alan H. Diercks, Alan Aderem, Alissa C. Rothchild

## Abstract

Alveolar macrophages (AMs) play a critical role during *Mycobacterium tuberculosis* (Mtb) infection as the first cells in the lung to encounter bacteria. We previously showed that AMs initially respond to Mtb *in vivo* by mounting a cell-protective, rather than pro-inflammatory response. However, the plasticity of the initial AM response was unknown. Here, we characterize how previous exposure to *mycobacterium*, either through subcutaneous vaccination with *Mycobacterium bovis* (scBCG) or through a contained Mtb infection (coMtb) that mimics aspects of concomitant immunity, impacts the initial response by AMs. We find that both scBCG and coMtb accelerate early innate cell activation and recruitment and generate a stronger pro-inflammatory response to Mtb *in vivo* by AMs. Within the lung environment, AMs from scBCG vaccinated mice mount a robust interferon-associated response, while AMs from coMtb mice produce a broader inflammatory response that is not dominated by Interferon Stimulated Genes. Using scRNAseq, we identify changes to the frequency and phenotype of airway-resident macrophages following *mycobacterium* exposure, with enrichment for both interferon-associated and pro-inflammatory populations of AMs. In contrast, minimal changes were found for airway-resident T cells and dendritic cells after exposures. *Ex vivo* stimulation of AMs with Pam3Cys, LPS and Mtb reveal that scBCG and coMtb exposures generate stronger interferon-associated responses to LPS and Mtb that are cell-intrinsic changes. However, AM profiles that were unique to each exposure modality following Mtb infection *in vivo* are dependent on the lung environment and do not emerge following *ex vivo* stimulation. Overall, our studies reveal significant and durable remodeling of AMs following exposure to *mycobacterium,* with evidence for both AM-intrinsic changes and contributions from the altered lung microenvironments. Comparisons between the scBCG and coMtb models highlight the plasticity of AMs in the airway and opportunities to target their function through vaccination or host-directed therapies.

**Author Summary:** Tuberculosis, a disease caused by the bacteria *Mycobacterium tuberculosis* (Mtb), claims around 1.6 million lives each year, making it one of the leading causes of death worldwide by an infectious agent. Based on principles of conventional immunological memory, prior exposure to either Mtb or *M. bovis* BCG leads to antigen-specific long-lasting changes to the adaptive immune response that can be effective at protecting against subsequent challenge. However, how these exposures may also impact the innate immune response is less understood. Alveolar macrophages are tissue-resident myeloid cells that play an important role during Mtb infection as innate immune sentinels in the lung and the first host cells to respond to infection. Here, we examined how prior *mycobacterium* exposure, either through BCG vaccination or a model of contained Mtb infection impacts the early innate response by alveolar macrophages. We find that prior exposure remodels the alveolar macrophage response to Mtb through both cell-intrinsic changes and signals that depend on the altered lung environment. These findings suggest that the early innate immune response could be targeted through vaccination or host-directed therapy and could complement existing strategies to enhance the host response to Mtb.

## Introduction

*Mycobacterium tuberculosis* (Mtb), the causative agent of Tuberculosis (TB), claimed more than 1.6 million lives in 2021. For the first time since 2005, the number of TB deaths worldwide is increasing (1, 2). These trends highlight the urgent need for new vaccine and therapeutic strategies. Traditionally, vaccine design has focused on generating a rapid, robust, and effective adaptive immune response. However, recent studies suggest that the innate immune system can undergo long-term changes in the form of trained immunity (3), which affect the outcome of infection and could function as important components of an effective TB vaccine (4, 5). Initial trained immunity studies focused on central trained immunity, long-term changes to hematopoietic stem cells that lead to functional changes in short-lived innate cell compartments (i.e., monocytes, NK cells, dendritic cells) (3). More recent studies have examined innate training in tissue-resident macrophages and demonstrated that these cells are also affected by prior exposures. Tissue-resident macrophages can respond to remote injury and inflammation (6), undergo long-term changes (3), and display altered responses to bacteria after pulmonary viral infection (7–9).

Lung resident alveolar macrophages (AMs) are the first cells to become infected with inhaled Mtb and engage a cell-protective response, mediated by the transcription factor Nrf2, that impedes their ability to effectively control bacterial growth (10, 11). In this study, we examined how prior mycobacterial exposure reprograms AMs and alters the overall innate response in the lung to aerosol challenge with Mtb. To evaluate the range of AM plasticity, we chose to compare the effects of subcutaneous BCG vaccination (scBCG) with those arising from a contained Mtb-infection (coMtb) model. BCG, a live-attenuated TB vaccine derived from *M. bovis* and typically given during infancy, provides protection against disseminated pediatric disease but has lower efficiency against adult pulmonary disease (12–14). In addition to enhancement of Mtb-specific adaptive responses, based on shared antigens, BCG vaccination also leads to changes in hematopoiesis and epigenetic reprogramming of myeloid cells in the bone marrow (15), early monocyte recruitment and Mtb dissemination (16), and innate activation of dendritic cells critical for T cell priming (17). Intranasal BCG vaccination protects against *Streptococcus pneumoniae* and induces long term activation of AMs (18). A recent study has shown that one mechanism by which BCG vaccination can elicit innate training effects on AMs, separate from alterations to the monocyte population, is through changes to the gut microbiome and microbial metabolites (19). BCG vaccination is also associated with trained immunity effects in humans (20–22), including well-described reductions in all-cause neonatal mortality and protection against bladder cancer (3, 23).

The coMtb model is generated by intradermal inoculation with virulent Mtb into the ears of mice and leads to a contained but persistent lymph node Mtb infection (24, 25). The model replicates observations in both humans and non-human primates (NHPs) that prior exposure to Mtb infection provides protection against subsequent exposure, through a form of concomitant immunity (26, 27). In previous studies, we found that coMtb leads to protection against challenge with aerosol Mtb infection and protects mice against heterologous challenges, including infection with *Listeria monocytogenes* and expansion of B16 melanoma cells, results which suggest there is substantial remodeling of innate immune responses (25). We previously found that AMs from coMtb mice mount a more inflammatory response to Mtb infection compared to AMs from control mice, and the enhancement in AM activation after infection, as measured by MHC II expression, was dependent on IFNγR signaling (25).

Here, we show that while both coMtb and scBCG protect against low dose Mtb aerosol challenge, they remodel the *in vivo* innate response in different ways. In AMs, scBCG elicits a very strong interferon response in AMs, while coMtb promotes a broader pro-inflammatory response that is less dominated by Interferon Stimulated Genes. Prior exposure to *mycobacterium* also remodels the frequency and phenotype of AM subsets in the lung prior to aerosol challenge and leads to significant changes in the early dynamics of the overall innate response. While changes in the AM responses that are unique to each exposure (scBCG, coMtb) depend on the lung environment, stronger interferon-associated responses following both LPS and Mtb stimulation *ex vivo* reveal cell-intrinsic changes.

## Results

### Prior mycobacterial exposure accelerates activation and innate cell recruitment associated with Mtb control

We first determined the earliest stage of infection when the immune response was altered by prior exposure to mycobacteria. Mice were vaccinated with scBCG or treated with coMtb, rested for 8 weeks, and then challenged with low-dose H37Rv aerosol infection. We measured both the cellularity and activation of innate immune cells in the lung at 10, 12 and 14 days following infection, the earliest timepoints when innate cells are known to be recruited (10, 11, 28). We observed a significant increase in MHC II Median Fluorescence Intensity (MFI) as early as day 10 for AMs from coMtb mice and day 12 for AMs from scBCG mice compared to controls (**Fig 1A, S1**). There were no significant differences in MHC II expression prior to challenge on day 0 (**Fig 1A**). There were also significant increases in the numbers of monocyte-derived macrophages (MDM), neutrophils (PMN), dendritic cells, and Ly6C^+^ CD11b^+^ monocytes by day 10 in coMtb mice compared to controls, with further increases by days 12 and 14 (**Fig 1B, Fig S1**). scBCG elicited similar increases in these populations starting at day 10, but the increases were not as robust or rapid as those observed in coMtb. Significant differences between scBCG and coMtb groups were found at d10, d12, d14 in MDM, d14 in PMN, d12, d14 in dendritic cells, and d14 in Ly6C^+^ CD11b^+^ monocytes (**Fig 1B**). While there were not significant differences in AM cell number between the three conditions, there was a modest drop in viability for both AMs from scBCG and coMtb mice by day 14 (**Fig 1C**).

**Figure 1:**
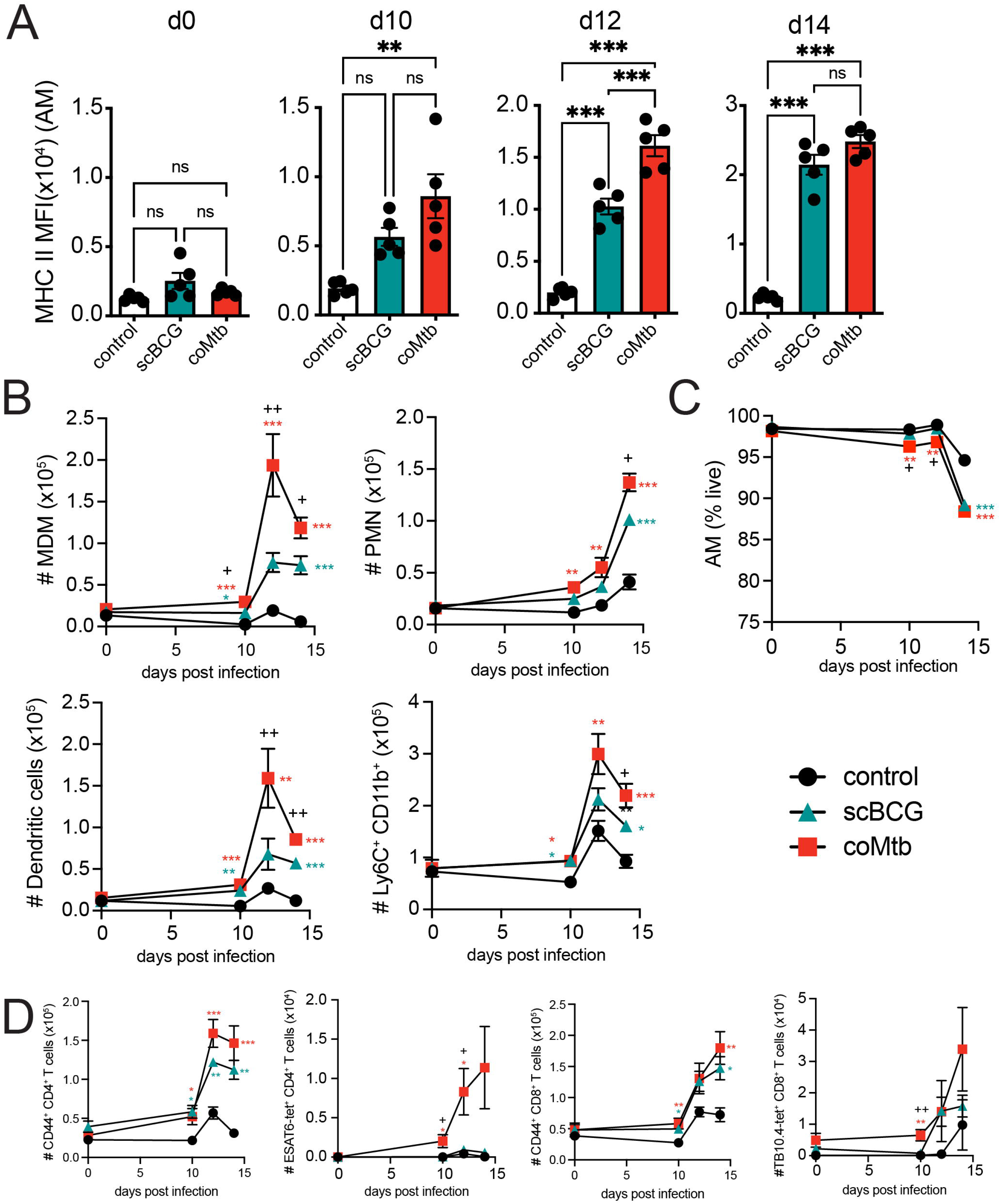
Prior exposure to *mycobacterium* leads to faster activation and innate cell recruitment following aerosol Mtb challenge. Control, scBCG, and coMtb mice, 8 weeks following exposure, challenged with standard low-dose H37Rv. Lungs collected on day 10, 12, and 14 post-infection. A) AM MHC II MFI. B) Total numbers of MDMs, PMN, DC, and Ly6C^+^CD11b^+^ monocytes. C) AM viability (% Zombie Violet-). D) Total numbers of CD44^+^ CD4^+^ T cells, ESAT6-tetramer^+^ CD4^+^ T cells, CD44^+^ CD8^+^ T cells, and TB10.4-tetramer^+^ CD8^+^ T cells. Mean +/-SEM, 5 mice per group, representative of 3 independent experiments. One-way ANOVA with Tukey post-test. * p< 0.05, **p< 0.01, ***p < 0.001. B, C) *, **, and *** scBCG or coMtb vs control; +, ++ scBCG vs coMtb.

In addition to early changes in innate cell activation and recruitment, we observed early recruitment of activated CD44^+^ CD4^+^ and CD8^+^ T cells in the lungs of both coMtb and scBCG mice starting at day 10 as well as TB antigen-specific T cells, ESAT6-tetramer^+^ CD4^+^ T cells and TB10.4-tetramer^+^ CD8^+^ T cells in coMtb mice starting at day 10 compared to controls and scBCG mice (**Fig 1D, Fig S1**). The differences in the recruitment of ESAT6-tetramer^+^ CD4^+^ T cells between scBCG and coMtb were expected, as the ESAT6 antigen is expressed by H37Rv but not by BCG.

We also evaluated whether these cell recruitment differences correlated with changes in bacterial burden. To compile CFU results from three independent experiments, each with slightly different bacterial growth (**Fig S2A**), we calculated a ΔCFU value that compared the bacterial burden of each sample to the average for the respective control based on timepoint, organ, and experiment. We found that both modalities generated a significant reduction in bacterial burden compared to controls in the lung, spleen, and lung-draining lymph node (LN) at day 14 and at day 28, as previously reported (16, 25, 29) (**Fig S2A-D)**. At day 10, we observed no difference in lung bacterial burden in scBCG or coMtb mice compared to controls and a small increase in coMtb mice over scBCG. The majority of control mice had undetectable bacteria in spleen and LN at this time. There was a significant reduction in bacterial burden in the lung by day 12 in coMtb but not scBCG mice and a significant reduction in CFU in the LN in both models compared to controls (**Fig S2B**). Our results demonstrate that prior mycobacteria exposure leads to accelerated innate cell activation and recruitment, alongside an increase in activated T cells, within the first two weeks of infection, with coMtb generating a faster and more robust response compared to scBCG. These early immune changes are associated with reductions in bacterial load in the lung, while differences in bacterial burden in the LN and spleen suggest delays in bacterial dissemination, which first appear in the LN at day 12 and then in the spleen at day 14 (**Fig S2A**).

### *Mycobacterium* exposure alters the *in vivo* alveolar macrophage response to Mtb infection

To examine the earliest response to Mtb, we measured the gene expression profiles of Mtb-infected AMs isolated by bronchoalveolar lavage and cell sorting, as previously described (10), 24 hours following aerosol challenge with high dose mEmerald-H37Rv (depositions: 4667, 4800) in scBCG-vaccinated mice and compared these measurements to previously generated profiles of AMs from control (unexposed) mice (10) and coMtb mice (25) (**Table S1**). As previously observed for the high dose infection, an average of 1.79% (range: 0.91-3.18%) of total isolated AMs were Mtb infected 24 hours after infection. Mtb-infection induced changes were measured by comparing gene expression between Mtb-infected AMs and respective naïve AMs for each of the three groups (control, scBCG, coMtb). Principal Component Analysis on Mtb-infected induced changes showed that each of the three conditions led to distinct expression changes (**Fig 2A**) and the majority of up-regulated Differentially Expressed Genes (DEG) (fold change > 2, FDR < 0.05) were unique to each condition (control: 151 unique/257 total genes, scBCG: 222/289, coMtb: 156/229) (**Fig 2B**). The divergence in the responses of Mtb-infected AMs from each of the 3 conditions was also reflected in the diversity in the Top 20 Canonical Pathways identified by Ingenuity Pathway Analysis (**Fig S3**).

**Figure 2:**
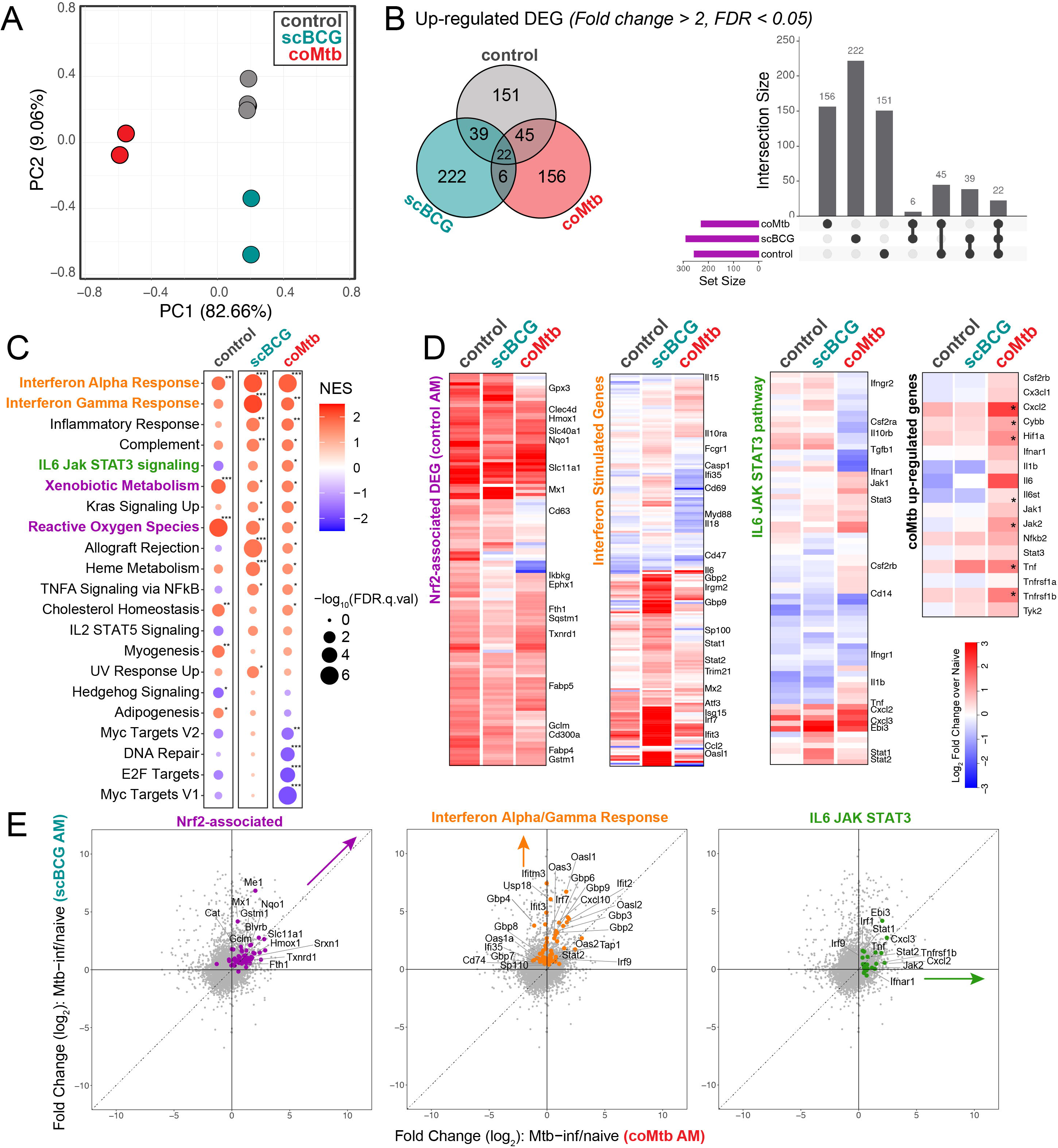
*Mycobacterium* exposure alters the alveolar macrophage transcriptional response to Mtb infection *in vivo*. Gene expression of Mtb-infected AMs 24 hours following high-dose mEmerald-H37Rv infection in mice previously exposed with scBCG or coMtb or controls (controls-reported in Rothchild et al, 2019 (10); coMtb-reported in Nemeth et al, 2020 (25)). A) Principal Component Analysis using DEG (|fold change| > 2, FDR < 0.05) in Mtb-infected AMs compared to respective naïve AMs (control, scBCG, or coMtb). B) Venn Diagram and Intersection plot of overlap in up-regulated DEG between the 3 conditions. C) Gene Set Enrichment Analysis of 50 Hallmark Pathways. Pathways shown have |NES| > 1.5 and FDR < 0.05 for at least one of the conditions. * FDR< 0.05, **FDR< 0.01, ***FDR< 0.001. D) Heatmap of 131 original *in vivo* DEG at 24 hours in Mtb-infected AM (*left*), Interferon Stimulated Genes, derived from macrophage response to IFNα (fold change >2, p-value < 0.01) Mostafavi et al, 2016 (30) (*center-left*), IL6 JAK STAT3 hallmark pathway (*center-right*) and selected coMtb signature genes (*right*, *FDR < 0.05, FC>2). E) Scatterplots depicting fold change (log_2_) for Mtb-infected AMs over naïve AMs for scBCG versus coMtb. Highlighted pathways: Nrf2-associated genes out of 131 original in vivo DEG (56 genes, purple), shared leading edge genes for scBCG Interferon Alpha Response and Interferon Gamma Response pathways (61 genes, orange), and leading edge genes for coMtb IL6 JAK STAT3 pathway (23 genes, green). Compiled from 4 independent experiments per condition for control, 2 independent experiments per condition for scBCG and coMtb.

To identify trends between groups, we performed Gene Set Enrichment Analysis using a set of 50 Hallmark Pathways. As we’ve shown previously, Mtb-infected AMs from control mice at 24 hours had strong enrichment for “Xenobiotic Metabolism” and “Reactive Oxygen Species” pathways, indicative of the Nrf2-associated cell-protective response (**Fig 2C**). While these two pathways were not among most enriched pathways in the exposed groups, Mtb-infected AMs from all groups upregulated genes associated with the 131 *in vivo* DEG that make up the cell-protective Nrf2-driven response at 24 hours (10) (**Fig 2D**). Expression profiles for Mtb-infected AMs from scBCG mice showed the strongest enrichment for “Interferon Alpha Response” and “Interferon Gamma Response” pathways, which contain many shared genes (**Fig 2C**). The strength of the interferon response was further highlighted by examining gene expression changes in a set of Interferon Stimulated Genes (ISGs) identified from macrophages responding to IFNα (fold change > 2, p-value < 0.01) (30) (**Fig 2D**). Expression profiles for Mtb-infected AMs from coMtb mice showed a weaker enrichment for interferon response pathways with fewer up-regulated ISGs compared to scBCG, and instead showed enrichment across a number of inflammatory pathways including “IL6 JAK STAT3 signaling” in comparison to the other groups (**Fig 2C, D**). A direct comparison between the gene expression patterns for AMs from scBCG versus coMtb mice could be visualized more readily by scatterplots highlighting either Nrf2-associated, Interferon Alpha and Gamma Response, or IL-6 JAK STAT3 pathway genes (**Fig 2E**).

In summary, mycobacteria exposures alter the initial *in vivo* response of AMs to Mtb infection 24 hours after challenge and remodel the AM response in distinct ways. AMs from scBCG vaccinated animals mount a strong interferon-associated response, while AMs from coMtb mice expression a more diverse inflammatory profile consisting of both interferon-associated genes as well as other pro-inflammatory genes, including those within the IL-6 JAK STAT3 pathway.

### *Mycobacterium* exposure modifies the baseline phenotype of alveolar macrophages in the airway

Although scBCG and coMtb exposures alter the AM responses to Mtb infection *in vivo*, transcriptional effects are not widely evident prior to infection as measured by bulk RNA-sequencing of naïve AMs from control, scBCG, or coMtb mice, including expression of innate receptors and adaptors (**Fig S4**). However, we posited that remodeling effects were likely not homogenous across the entire AM population and that small heterogenous changes to baseline profiles might be detectable using a single cell approach. We therefore analyzed pooled BAL samples taken from 10 age- and sex-matched mice from each of the three conditions (control, scBCG, coMtb) eight weeks following mycobacteria exposure by single cell RNA-sequencing (scRNAseq). Gross cellularity was unaffected by mycobacterial exposure as measured by flow cytometry analysis of common lineage markers with AMs the dominant hematopoietic cell type (57.4-85.8% of CD45^+^ live cells), followed by lymphocytes (5.26-22.7% of CD45^+^ live cells) with small contributions from other innate cell populations (**Fig S5**).

Six samples, with an average of 2,709 cells per sample (range: 2,117-4,232), were analyzed together for a total of 17,788 genes detected. The most prominent expression cluster mapped to an AM profile, with smaller clusters mapping to T and B lymphocytes, dendritic cells, and neutrophils (**Fig 3A**). All cells that mapped to a macrophage profile were extracted and reclustered into 11 macrophage subclusters (**Fig 3B, C**). All but two of the macrophage subclusters (clusters 6 and 8) expressed AM lineage markers (*Siglecf, Mertk, Fcgr1* (CD64), *Lyz2* (LysM), and *Itgax* (CD11c) and had low expression of *Itgam* (CD11b) (**Fig 3D**). Cluster 6 showed high *Itgam* and *Lyz2* expression and lower *Siglecf* expression, likely representing a small monocyte-derived macrophage population in the airway, while cluster 8 displayed high *Lyz2* expression, low expression for other AM markers, and expression of *S*ftpa1 and *Wfdc2* (**Table S2**), genes most commonly expressed by pulmonary epithelial cells, suggesting that this cluster represents a small population of epithelial cells,

**Figure 3:**
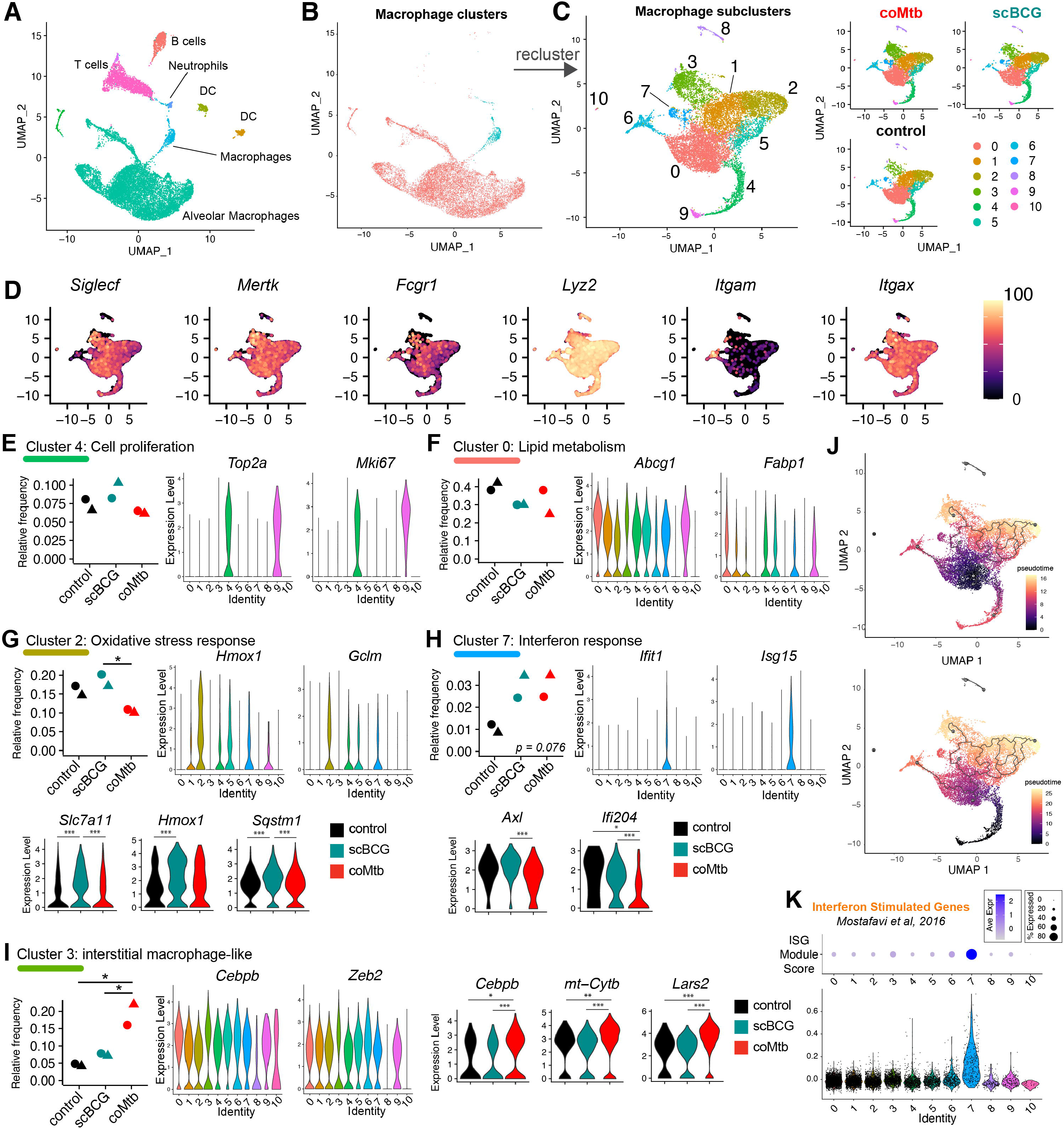
*Mycobacterium* exposure modifies the alveolar macrophage phenotype in the airway pre-challenge. Single-cell RNA-sequencing of BAL samples from control, scBCG, and coMtb mice. A) Compiled scRNAseq data for all BAL samples, highlighted by major clusters, annotated based on closest Immgen sample match. B) Highlighting of the two clusters used for macrophage subcluster analysis. C) The 12 clusters generated by the macrophage subcluster analysis, separated by condition. D) Expression of major macrophage-specific markers: *Siglecf, Mertk, Fcgr1, Lyz2, Itgam* (CD11b), and *Itgax* (CD11c). E-I) Relative frequency of each macrophage subcluster by condition. (*violin plots by cluster*) Expression level of representative genes distinguished by that cluster compared to other clusters. One-way ANOVA with Tukey post-test, * p< 0.05. (*3-way violin plots by condition*) Differentially expressed genes within Clusters 2, 7, and 3 between control vs scBCG vs coMtb samples. Wilcoxon Rank Sum Test, Bonferroni adjusted p-value. ***adj-p < 0.001, **adj-p < 0.01, *adj-p < 0.05. J) Pseudotime analysis (Monocle3) with starting node at the largest cluster in control, Cluster 0 (*top*) and at the cluster of proliferating cells, Cluster 4,9 (*bottom*). K) ISG Module Score by cluster. Module derived from macrophage response to IFNα (fold change > 2, p-value < 0.01) (Mostafavi et al, 2016). Data is compiled from two independent experiments (circle, triangle) with 3 conditions each for a total of 6 samples.

To interpret the various expression subclusters, we identified the genes that most distinguished each cluster from the others (**Fig S6, Table S2**). As has been reported by other groups (31, 32), a small proportion of the AMs in two clusters (Clusters 4, 9) with high expression of cell cycle genes (i.e., *Top2a, Mki67*), indicative of cell proliferation (**Fig 3E, Table S2**). Cluster 0 formed the most abundant macrophage cluster with high expression of lipid metabolism genes (i.e., *Abcg1, Fabp1*) (**Fig 3F, Table S2**). Cluster 2 was significantly increased in relative frequency for scBCG samples compared to coMtb (p = 0.032, *One-way ANOVA with Tukey post-test*) and associated with oxidative stress response genes (*Hmox1, Gclm*). Several Cluster 2 associated genes, *Slc7a11, Hmox1,* and *Sqstm1* also had higher overall expression level in scBCG samples compared to either control or coMtb **(Fig 3G, Table S2**). Cluster 7 was the only cluster with an increase in relative frequency trending for both scBCG and coMtb (p = 0.076, *One-way ANOVA*). Cells in this cluster had high expression of Interferon Stimulated Genes (*Ifit1, Isg15*) and within this cluster, cells from scBCG samples had higher expression of *Axl* and *Ifi204* than cells from coMtb samples. (**Fig 3H, Table S2**). Cluster 3 had significantly higher relative frequency for coMtb samples compared to control and scBCG samples (p = 0.021, 0.039, respectively, *One-way ANOVA with Tukey post-test*) and was distinguished by expression of macrophage-associated transcription factors (*Cebpb*, *Zeb2*, *Bhlhe40*) (33, 34), mitochondrial oxidative phosphorylation (*mt-Co1, mt-Cytb, mt-Nd2*), chromatin remodeling (*Ankrd11, Baz1a*), and immune signaling including the CARD9 complex (*Malt1, Bcl10, Prkcd*) (**Fig 3I, S7, Table S2**). This expression profile closely matches a subcluster of AMs previously described by Pisu et al, as an “interstitial macrophage-like” AM population (labeled “AM_2”) that expanded in relative frequency in lung samples 3 weeks following low-dose H37Rv infection (31). Relative expression level for *Cebpb, Mt-Cyb*, and *Lars2* within Cluster 3 was higher for cells from coMtb samples compared to either control or scBCG samples.

Interestingly, Cluster 2 (higher relative frequency in scBCG) and Cluster 3 (higher relative frequency in coMtb) represent divergent endpoints of a pseudotime plot generated by a trajectory inference analysis, regardless of whether the starting point is the most abundant cluster in the control group (Cluster 0) (**Fig 3J, *top***) or the cluster of proliferating cells (Cluster 4) (**Fig 3J, *bottom***). This result suggests that scBCG and coMtb may drive AM phenotypes in divergent directions and indicates that AM responses can be remodeled into more than one flavor, rather than only a binary “on/off” state.

To further investigate whether a sub-cluster of AMs might be responsible for the increased enrichment for Interferon Alpha/Gamma Response pathways in the *in vivo* Mtb response in scBCG and coMtb mice, we scored each cluster based on the ISG gene module, previously used in **Fig 2D**. As expected, we observed that only Cluster 7 showed strong enrichment for ISGs, which was trending up in frequency for both scBCG and coMtb samples (**Fig 3K**).

In summary, scRNAseq analysis of macrophages isolated by BAL demonstrate that *mycobacterium* exposure leads to subtle changes in small minority subsets of AMs in the airway, including subsets associated with interferon responses and an interstitial macrophage phenotype, while leaving the largest subsets of AMs unchanged in frequency or gene expression. We hypothesize that these small changes in baseline profiles may be sufficient to drive the more substantial changes observed in the AM Mtb response *in vivo*, as described in **Figure 2**.

### *Mycobacterium* exposure has minimal impact on T cell populations in the airway

While AMs are the dominant immune cell type in the airway, other cell populations make up an average of 18.4% of the cells within the BAL in controls (range: 10.4-26.3%) and 31.3% in exposed groups (range: 14.0-48.8%). To examine how mycobacteria exposure influenced other cells in the airway, we focused on T cells and dendritic cells (DCs) which have two of the highest relative frequencies after AMs (**Fig 4A, B**). T cells and DCs were each combined from two original clusters each. Neither population showed a statistically significant difference in relative frequency (**Fig 4B**). To examine qualitative changes in the T cell population in greater detail, we next reclustered the T cells, resulting in 7 T cell clusters. We manually annotated each of the clusters based on the most closely matched Immgen profiles and the expression of key lineage specific markers (**Fig 4A-C, Fig S8**). We focused on the 5 most abundant T cell subclusters (Clusters 0-4). While we observed subtle shifts in the relative frequency of each group, none reached statistical significance. Cluster 0, the most abundant cluster, had an expression profile most consistent with γδ T cells, including expression of *Cd3e* with low to nil *Cd4* and *Cd8a* and some expression of *Zbtb16* (PLZF) and *Tmem176a*, an ion channel regulated by RORγt and reported to be expressed by lung γδ T cells (35, 36) (**Fig 4D-F, S8**). Cluster 1 matched a profile for effector CD4^+^ T cells (**Fig 4D-F, S8**), and Cluster 2 matched a profile for naïve CD8^+^ T cells (**Fig 4D-F, S8**). Cluster 3 had a profile consistent with effector memory/resident memory CD8^+^ T cells (T_EM/RM_) (**Fig 4D-F, Fig S7**) and Cluster 4 had a profile consistent with NK cells. Overall, there were no significant changes in the relative frequency of T cell or NK subclusters, despite detection of a number of different lymphocyte subsets in the airway.

**Figure 4:**
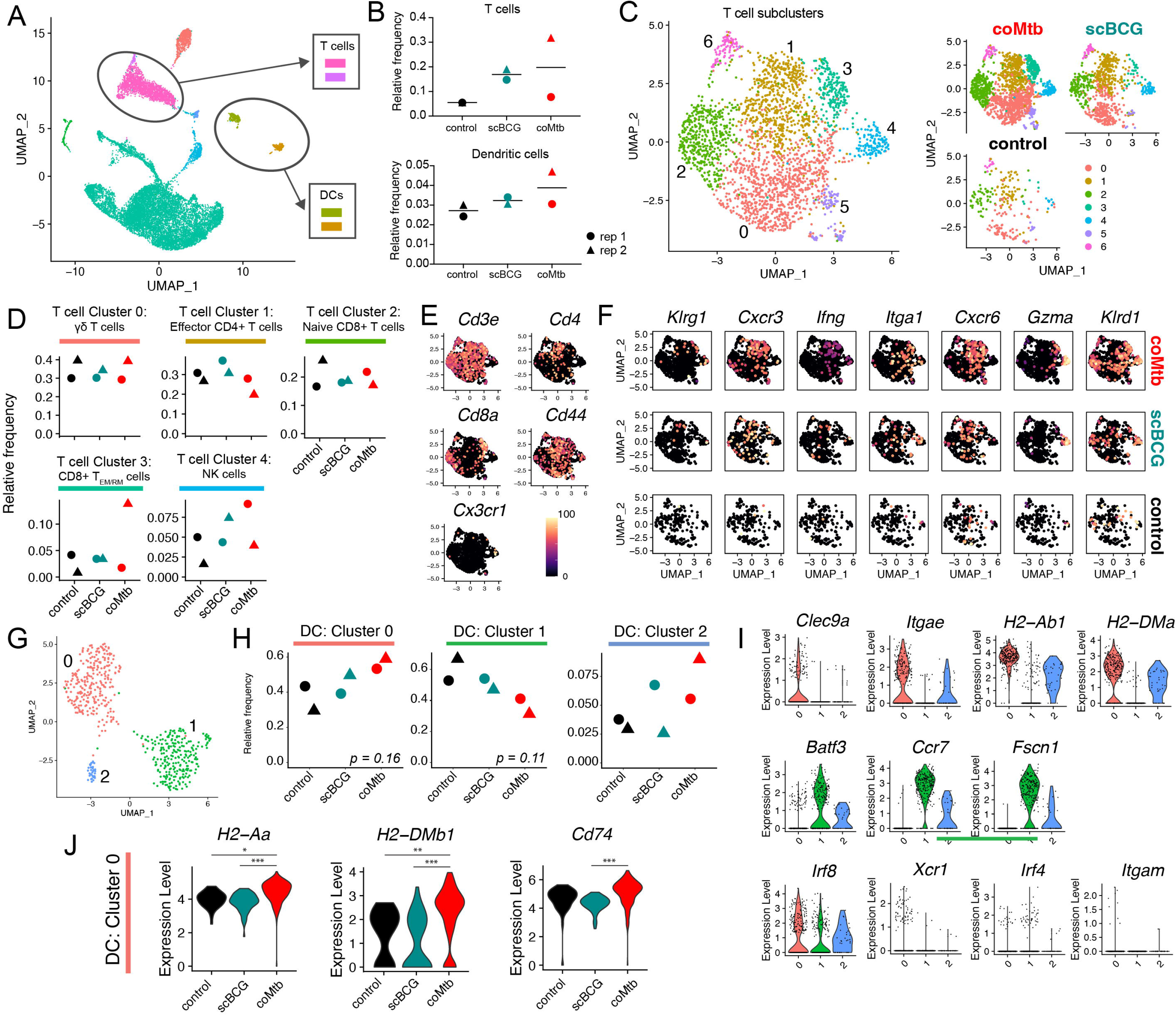
Airway T cell and dendritic cell profiles following *mycobacterium* exposure. Single-cell RNA-sequencing of BAL samples from control, scBCG, and coMtb mice. A) Compiled scRNAseq data for all BAL samples, with T cell and dendritic cell clusters highlighted. B) Relative frequency of T cells and DCs. C-F) T cell subcluster analysis. C) T cell subclusters compiled and split by condition. Annotations made following Immgen profile matches and manual marker inspection. D) Relative frequency of Clusters 0-4 for each condition. E) UMAP gene expression plot for general T cell markers. F) UMAP gene expression plot cluster-specific markers split by condition. G-I) Dendritic cell subcluster analysis. G) Dendritic cell subcluster, colored by each of 3 different clusters. H) Relative frequency of Clusters 0-2 for each condition. I) Violin plots for cluster-specific markers and genes of interest. J) Differentially expressed genes in Cluster 0 split by condition. *adj-p < 0.05, **adj-p < 0 .01, ***adj-p < 0.001, Wilcoxon Rank Sum Test, Bonferroni adjusted p-values. Data is compiled from two independent experiments with 3 conditions each for a total of 6 samples.

### *Mycobacterium* exposure modifies the dendritic cell airway landscape

Re-clustering of DCs yielded 2 major clusters (Cluster 0, 1) and 1 minor cluster (Cluster 2), which had a mixed phenotype with expression of genes from both major clusters (**Fig 4G**). Cells in Cluster 0 had high expression of *Clec9a, Itgae* (CD103), and MHC II genes (*H2-Ab1, H2-DMa*) consistent with an expression profile of lung CD103^+^ cDCs (37) (**Fig 4I**), while Cluster 1 had higher expression for *Batf3, Ccr7*, and *Fscn1*. All three of the clusters had high *Irf8* expression and low expression of *Xcr1*, *Irf4*, and *Itgam* (CD11b) (**Fig 4I**). While the coMtb samples trended higher in relative frequency for Cluster 0 and low for Cluster 1, compared to the control or scBCG samples, these differences did not meet statistical significance (One-way ANOVA with Tukey post-test, p = 0.16, p = 0.11), likely due to the limit in statistical power with only 2 replicates (**Fig 4H**). However, it was notable that for cells within Cluster 0, there was a significantly higher expression level for MHC II genes (*H2-Aa, H2-DMb1*, and *Cd74*) for coMtb cells compared to control or scBCG cells (**Fig 4J**). This suggests that coMtb might be able to elicit more mature or activated DCs in the airway. Overall, scRNAseq analysis shows that *mycobacterium* exposure leads to minimal changes in T cell and dendritic cell populations in the airway, although we hypothesize that small changes in DC maturation/activation could have important impacts on adaptive immune priming dynamics after subsequent aerosol infection.

### Cell-intrinsic remodeling of alveolar macrophages following *mycobacterium* exposure licenses an interferon response *in vitro*

Analysis of the AM response to Mtb *in vivo* demonstrates that the very earliest immune response to Mtb is altered by previous *mycobacterium* exposure. However, one limitation to this approach is the inability to discern whether changes to AMs are cell-intrinsic or dependent on the altered tissue environment, especially the presence of Mtb-specific T cells. Therefore, to determine whether mycobacteria exposure induces cell-intrinsic changes to AMs, we isolated AMs from control, scBCG, and coMtb mice, stimulated them *ex vivo* with LPS, Pam3Cys, or H37Rv, and measured their transcriptional profile 6 hours later (**Fig 5A**). First, PAMP-specific trends were notable. AMs from coMtb and scBCG mice showed distinct responses from AMs from control mice following LPS and H37Rv stimulation, but only minimal changes following Pam3Cys stimulation, with no obvious changes in innate receptor or adaptor expression to explain the PAMP-specific differences (**Fig 5B, S9, Table S3**). Second, as we have previously reported, Mtb-infected AMs did not strongly up-regulate Nrf2-associated genes *ex vivo* (**Fig 5C**). Third, when we looked at the gene sets that distinguished the *in vivo* AM response between scBCG and coMtb mice, “Interferon Alpha/Gamma Response” and “IL6 JAK STAT3” (**Fig 2E**), we found that the differences between exposure modalities was diminished *ex vivo*, suggesting contribution of the lung environment to the quality of the response (**Fig 5C**). Using Gene Set Enrichment Analysis, we identified “Interferon Gamma Response”, “Interferon Alpha Response”, “TNFa signaling via NF-kB”, and ”Inflammatory Response” pathways as the most strongly enriched for LPS and H37Rv responses from scBCG and coMtb AM (**Fig 5D)**. To assess whether the cell-intrinsic changes observed were long-lasting, we compared the responses of AMs at 8 or 23 weeks following scBCG vaccination by RT-qPCR. Increases in gene expression were as robust or even enhanced 23 weeks following exposure compared to 8 weeks, suggesting that exposure-induced changes to AMs are relatively long-lived (**Fig S10**). To validate whether changes in gene expression were reflected at the protein level, we sought to develop a flow cytometry-based assay to assess AM-specific responses. Primary AMs were stimulated with LPS for 20 hours and both MHC II and TNF expression were measured by flow cytometry **(Fig 5E**). We found that AM from coMtb mice had significantly higher MHC II expression than controls and a similar pattern was seen for scBCG AM in 1 of 2 experiments (**Fig 5F**). AMs from coMtb mice also showed a significant increase in TNF expression in 1 of 2 experiments (**Fig 5G**).

**Figure 5:**
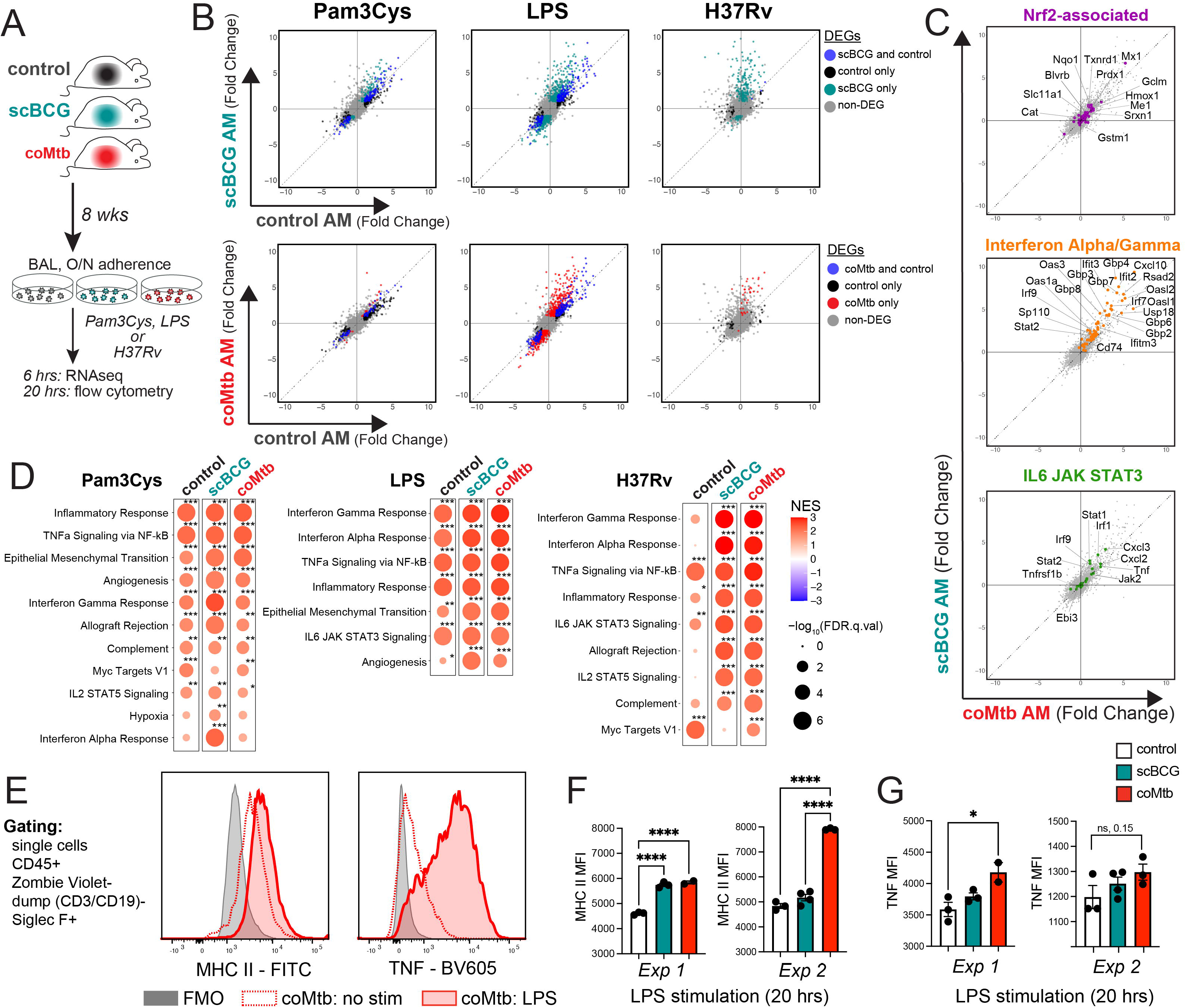
Cell-intrinsic remodeling of alveolar macrophages following *mycobacterium* exposure. A) AM isolation 8 weeks following scBCG or coMtb exposure. AMs were stimulated with Pam3Cys (10 ng/ml), LPS (10 ng/ml), and H37Rv (effective MOI ∼2:1) for 6 hours (RNA-seq) or 20 hours (flow cytometry). B) Scatterplots, log_2_ fold change gene expression for stimulated to unstimulated AMs for each condition (control, scBCG, coMtb). Differentially expressed genes (DEG) are highlighted for one or both conditions (|Fold change| > 2, FDR < 0.05 for Pam3Cys and LPS; |Fold change| > 2, FDR < 0.2 for H37Rv). C) Scatterplots, log_2_ fold change gene expression for H37Rv-stimulated to unstimulated scBCG versus coMtb AMs. Genes highlighted derived from gene sets in Fig 2E. Nrf2-associated genes (56 genes, purple), Interferon Alpha/ Gamma Response (61 genes, orange), and IL6 JAK STAT3 (23 genes, green). D) Gene Set Enrichment Analysis results for 50 HALLMARK pathways. Pathways shown have NES > 1.5 and FDR < 0.05 for at least one of the conditions. * FDR < 0.05, ** FDR < 0.01, *** FDR < 0.001. E) Gating strategy and MHC II and TNF histograms for coMtb AM, no stimulation versus LPS. F-G) MHC II and TNF MFI in control, scBCG, and coMtb AM after 20 hours of LPS stimulation. * p < 0.05, ** p < 0.01, *** p < 0.001, One-way ANOVA with Tukey post-test.

Because the “Interferon Alpha Response” and “Interferon Gamma Response” pathways were most highly enriched for the H37Rv stimulation following mycobacteria exposure, we decided to further investigate the Interferon-associated response (30). We categorized the macrophage response to H37Rv stimulation as “IFN-dependent” or “IFN-independent” based on gene expression of WT and IFNAR^−/−^ bone marrow derived macrophages (BMDMs) following H37Rv infection (see methods section) (**Table S4**) (38). Expression of IFN-dependent genes was minimally induced in control AMs but strongly up-regulated in AMs from *mycobacterium* exposed mice, as measured by the GSEA normalized enrichment score (NES) (**Fig 6A, *left***). In contrast, expression of IFN-independent genes was modestly upregulated in control AMs and only slightly altered by *mycobacterium* exposure (**Fig 6A, *right***). When we applied these two gene sets to the *in vivo* response profiles described in Figure 2, we observe that Mtb-infected AM from scBCG mice up-regulate the IFN-dependent response *in vivo*, suggesting that the licensing of the IFN-dependent response plays a role *in vivo* following BCG vaccination (**Fig 6B**). The difference between the *in vitro* and *in vivo* response for AM from coMtb mice points to an additional contribution of the lung environment.

**Figure 6:**
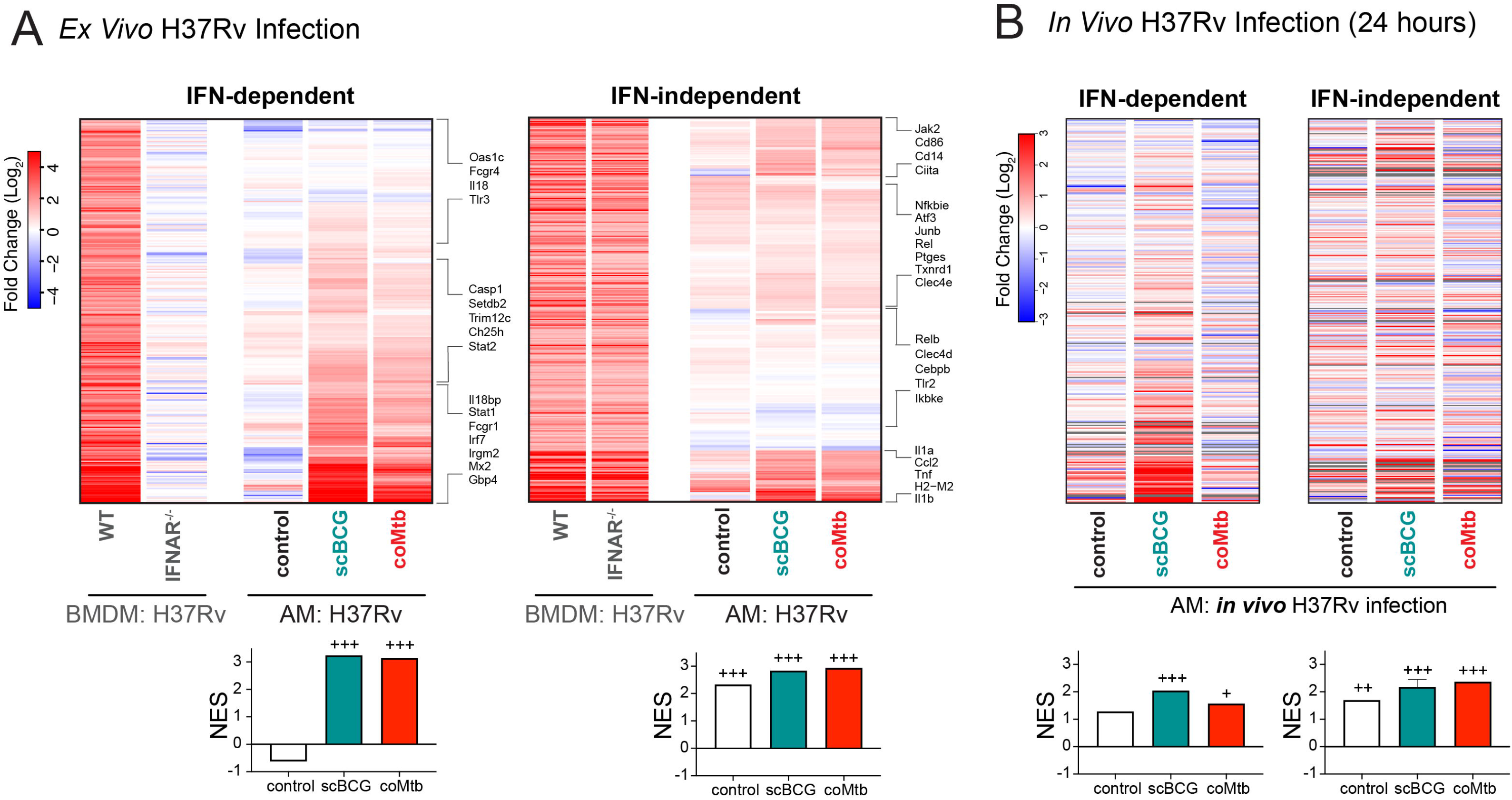
*Mycobacterium* exposure licenses an interferon-dependent response to H37Rv by alveolar macrophages. A) Gene expression for control, scBCG, and coMtb AMs, 6 hour H37Rv infection *ex vivo*, log_2_ fold change (Mtb-infected/uninfected). IFN-dependent genes (339 total) and IFN-independent genes (352 total) based on WT vs IFNAR^−/−^ BMDM bulk RNA-seq dataset (Olson et al, 2021) (see Methods section). B) Gene expression for control, scBCG, and coMtb AMs, 24 hour H37Rv aerosol infection *in vivo*, Mtb-infected sorted, log_2_ fold change (Mtb-infected/uninfected) for the same IFN-dependent and IFN-independent gene sets in (A). Grey bars indicate N.D. Normalized Enrichment Score (NES) calculated by GSEA for two data sets alongside Hallmark Pathways. + FDR < 0.05, ++ FDR< 0.01, +++ FDR < 0.001.

These results demonstrate that prior *mycobacterium* exposure leads to cell-intrinsic changes in AMs that license an enhancement of IFN-dependent responses to Mtb that are retained *in vitro*, while qualitative differences in the response between scBCG and coMtb *in vivo* depend on signals from the lung environment.

## Discussion

Here we describe remodeling of AMs, long-lived airway-resident innate cells, following two modalities of *mycobacterium* exposure, scBCG vaccination and coMtb, a model of contained Mtb infection. AMs are the first cells to be productively infected in the lung following aerosol Mtb infection (10, 11). We previously showed that AMs initially respond to Mtb infection with a cell-protective, Nrf2-driven program that is detrimental to early host control (10), suggesting that the lack of a robust response by AMs prevents effective host control early on. In line with this model, others have shown that depletion of AMs or strategies that “bypass” AMs including directly injecting antigen-primed DCs or activating DCs accelerates the immune response and reduces bacterial burden (17, 39, 40). However, how vaccination or prior exposures impact the initial response of AMs and whether there are therapeutic strategies that would enhance their initial response to infection have not been well studied (41).

Most studies examining the impacts of prior exposure to either Mtb or other species of mycobacteria, including BCG vaccination, have focused on the durable changes to the adaptive immune response that are antigen-specific. In contrast, we focused on changes to tissue-resident innate cells and their responses at the earliest stages of infection (≤ 14 days). Along with early changes to the T cell response, both scBCG and coMtb accelerate innate cell activation and immune cell recruitment in the first 10-14 days following Mtb aerosol infection, and even the very initial AM response to Mtb, within the first 24 hours of infection, is remodeled following exposure to *mycobacterium*. The durable changes observed fit with a number of recent studies, which have uncovered either enhanced AM antimicrobial phenotypes (7–9) or impaired responses (42, 43) following viral infection. Other studies have identified long-lasting changes to AMs following intranasal immunization of either adenoviral-based or inactivated whole cell vaccines (18, 44, 45).

We observe that the most robust cell-intrinsic changes to AM responses following scBCG or coMtb are in IFN-dependent genes (Fig 6) and ISGs (Fig 2D), suggesting a critical role for interferon signaling in the changes to the early innate response in the lung during infection. Notably, this finding is not limited to the murine model. BAL from NHPs following IV, ID, or aerosol BCG vaccination similarly show AMs enriched for Interferon Gamma Response pathway genes (46). AMs can respond to both Type I (IFNα/β) and Type II Interferons (IFNγ) and it is not possible to distinguish between responses to IFNα/β and IFNγ based on transcriptional analysis alone. The presence of live bacteria within both scBCG and coMtb models limits system-wide perturbations, such as T cell depletion or anti-IFNγ blockade, which would reverse containment (24). For this reason, we have not been able to directly test how interferon signals derived from scBCG or coMtb remodel AMs in a cell-autonomous manner, but we envision future studies to examine the specific effects of individual cytokines on AM remodeling.

IFNγ is a likely candidate to contribute to AM remodeling following *mycobacterium* exposure. IFNγ is required for the generation of trained immunity in bone marrow-derived myeloid cells following IV BCG vaccination (15, 47). While IV and aerosol H37Rv infection was found to induce Type I IFNs and reduce myelopoiesis (47), we previously found that coMtb, in which Mtb is contained within the ear-draining lymph node, leads to low-level systemic cytokinemia, including IFNγ production. Using WT:Ifngr1^−/−^ mixed bone marrow chimeras, we showed that IFNγ signaling was responsible for monocyte and AM activation following establishment of coMtb (25). Additionally, several reports have identified T cell-derived IFNγ as critical for altering AM function, although the immunological outcome varies substantially based on the context. In one study, T cell-derived IFNγ following adenoviral infection leads to AM activation, innate training and protection from *S. pneumoniae* (8), while in another study influenza-induced T cell-derived IFNγ leads to AM dysfunction and impaired clearance of *S. pneumoniae* (43). A study of 88 SARS-CoV-2 patients identified AMs and T cell-derived IFNγ as part of a positive feedback loop in the airway (48). In contrast, type I IFN signatures are associated with active TB or TB disease progression in both humans and non-human primates (49–51). Host perturbations such as treatment with poly I:C or viral co-infection that induce type I IFN lead to worsened disease (52, 53), type I IFN has been shown to block production of IL-1β in myeloid cells during Mtb infection (54), and type I IFN drives mitochondrial stress and metabolic dysfunction in Mtb infected macrophages (38).

We note that the two modalities tested here consist of different mycobacterial species, different doses, and different routes. We expect that all three of these factors likely contribute to the quality of AM remodeling. For example, they could be important for the location, timing, and amount of IFNγ that AMs are exposed to. While an in-depth examination of each of these factors is beyond the scope of this study, the side-by-side comparison of the two different exposure models, scBCG and coMtb, allows us to examine the plasticity of AM phenotypes and the impact of the local and/or systemic environments leading to different responses. In particular, the disappearance of the modality-specific signatures following *ex vivo* isolation hint at a critical contribution of the altered lung microenvironments in AM remodeling. The fact that AMs can be remodeled into more than one state suggests additional complexity in innate immune features that has not yet been fully explored. Heterogeneity in myeloid reprogramming is not limited to the murine model and has also been observed in human monocytes (55).

Several studies have recently described innate-adaptive interactions within the airway that are thought to impact infection dynamics (46, 48). We note that in these models we observe innate cell activation and recruitment occurring at the same time as T cell activation and recruitment, and that these events are likely promoting one another. We are particularly intrigued by the changes in AM MHC class II expression that we observed *in vivo* during the first two weeks of infection (**Fig 1A**) and following *ex vivo* stimulation (**Fig 5F**). AMs are considered to be poor antigen presenters, relative to other myeloid subsets, yet the faster Mtb-specific T cells are recruited to the lung, the more likely AMs will be to serve as primary T cell targets (37, 56-60). Our results suggest that enhancement of AM antigen presentation could be one innate mechanism that could be targeted to complement and synergize with the adaptive immune response during infection.

There are many other remaining questions. While we identify both cell-intrinsic changes and changes dependent on the lung environment, we do not yet know whether the cell-intrinsic changes are retained long-term in the absence of environmental cues. We do not know the durability of the changes, both cell-intrinsic and environment-dependent, and whether they are mediated by epigenetic effects. Our longest experiment showed retention of cell-intrinsic changes to AMs after 23 weeks. In Nemeth et al, we showed that antibiotic treatment lessened the protection mediated by coMtb, suggesting that ongoing replication is a key part of host protection (25). AM remodeling is retained 8 weeks or longer after the initial exposures, a timepoint when there is little to no detectable mycobacteria in the lung, ruling out a requirement for local ongoing bacterial replication in AM remodeling, although systemic signals derived from remote bacterial replication may still play a role. We also performed several of these studies with intravenous BCG vaccination (ivBCG), which in the mouse model leads to more sustained bacterial replication than scBCG (data not shown) (61). While we observed similar remodeling to AMs in the ivBCG model, these were not different in quality to those of scBCG vaccination, despite the major differences in bacterial replication and far greater T cell recruitment to the airway, suggesting that these changes are not required for AM remodeling.

There is still much unknown about the signals that drive reprogramming of tissue-resident innate cells. Ideally, vaccines would be designed to leverage these signals in order to promote the most effective interactions between innate and adaptive responses. Identifying the ways that AMs are reprogrammed by inflammatory signals and the effects of their changed phenotypes on the early stages of infection will help to improve future vaccines or host-directed therapies.

## Materials and Methods

### Mice

C57BL/6 mice were purchased from Jackson Laboratories (Bar Harbor, ME). Mice were housed and maintained in specific pathogen-free conditions at Seattle Children’s Research Institute and University of Massachusetts Amherst and experiments were performed in compliance with the respective Institutional Animal Care and Use Committees. 6-12 week old male and female mice were used for all experiments, except for RNA-sequencing, which used only female mice for uniformity. Mice infected with Mtb were housed in Biosafety Level 3 facilities in Animal Biohazard Containment Suites.

### Mycobacteria exposure models: BCG immunization and establishment of coMtb

BCG-Pasteur was cultured in Middlebrook 7H9 broth at 37°C to an OD of 0.1–0.3. Bacteria was diluted in PBS and 1 x 10^6^ CFU in 200 ml was injected SC. Intradermal infections to establish coMtb were performed as formerly described (24), with some modifications as detailed previously (25). Briefly, 10,000 CFU of Mtb (H37Rv) in logarithmic phase growth were injected intradermally into the ear in 10 μL PBS using a 10 μL Hamilton Syringe, following anesthesia with ketamine/xylazine.

### *M. tuberculosis* Aerosol Infections and Lung Mononuclear Cell Isolation

Aerosol infections were performed with wildtype H37Rv, including some transformed with an mEmerald reporter pMV261 plasmid, generously provided by Dr. Chris Sassetti and Christina Baer (University of Massachusetts Medical School, Worcester, MA). For both standard (∼100 CFU) and high dose (1,473-4,800 CFU) infections, mice were enclosed in an aerosol infection chamber (Glas-Col) and frozen stocks of bacteria were thawed and placed inside the associated nebulizer. To determine the infectious dose, three mice in each infection were sacrificed one day later and lung homogenates were plated onto 7H10 plates for CFU enumeration. High dose challenge and sorting of Mtb-infected AM was performed 4 weeks following scBCG vaccination and 2 weeks following coMtb vaccination as previously described (62). All other analysis was performed 8 weeks following mycobacterium exposures.

### Lung Single Cell Suspensions

At each time point, lungs were removed, and single-cell suspensions of lung mononuclear cells were prepared by Liberase Blendzyme 3 (70 ug/ml, Roche) digestion containing DNaseI (30 µg/ml; Sigma-Aldrich) for 30 mins at 37°C and mechanical disruption using a gentleMACS dissociator (Miltenyi Biotec), followed by filtering through a 70 µM cell strainer. Cells were resuspended in FACS buffer (PBS, 1% FBS, and 0.1% NaN_3_) prior to staining for flow cytometry. For bacterial enumeration, lungs were processed in 0.05% Tween-80 in PBS using a gentleMACS dissociator (Miltenyi Biotec) and were plated onto 7H10 plates for CFU enumeration. ΔCFU (log) was calculated as follows: ΔCFU = log((sample CFU)/(average control CFU*). *For respective experiment, timepoint, and organ. A ΔCFU value of −1 corresponds to a 10-fold reduction in CFU for the sample, compared to the control. Similarly, a ΔCFU value of 1 corresponds to a 10-fold increase in CFU.

### Alveolar Macrophage Isolation

AMs for cell sorting, bulk RNA-sequencing, single cell RNA-sequencing, and ex vivo stimulation were collected by bronchoalveolar lavage (BAL). BAL was performed by exposing the trachea of euthanized mice, puncturing the trachea with Vannas Micro Scissors (VWR) and injecting 1 mL PBS using a 20G-1” IV catheter (McKesson) connected to a 1 mL syringe. The PBS was flushed into the lung and then aspirated three times and the recovered fluid was placed in a 15mL tube on ice. The wash was repeated 3 additional times. Cells were filtered and spun down. For antibody staining, cells were suspended in FACS buffer. For cell culture, cells were plated at a density of 5 x 10^4^ cells/well (96-well plate) in complete RPMI (RPMI plus FBS (10%, VWR), L-glutamine (2mM, Invitrogen), and Penicillin-Streptomycin (100 U/ml; Invitrogen) and allowed to adhere overnight in a 37°C humidified incubator (5% CO_2_). Media with antibiotics were washed out prior to infection with *M. tuberculosis*.

### Cell Sorting and Flow Cytometry

Fc receptors were blocked with anti-CD16/32 (2.4G2, BD Pharmingen). Cell viability was assessed using Zombie Violet dye (Biolegend). Cells were suspended in 1X PBS (pH 7.4) containing 0.01% NaN_3_ and 1% fetal bovine serum (i.e., FACS buffer). Surface staining, performed at 4 degrees for 20 minutes, included antibodies specific for murine: Siglec F (E50-2440, BD Pharmingen), CD11b (M1/70), CD64 (X54-5/7.1), CD45 (104), CD3 (17A2, eBiosciences), CD19 (1D3, eBiosciences), CD11c (N418), I-A/I-E (M5/114.15.2), Ly6G (1A8), Ly6C (HK1.4), TNF (MP6-XT22). For ICS, Brefeldin A was added for duration of LPS stimulation. Cyto-Fast Fix/Perm and Cyto-Fast Perm Wash reagents were used for intracellular staining. Reagents are from Biolegend unless otherwise noted. MHC class II tetramers ESAT-6 (I-A(b) 4–17, sequence: QQWNFAGIEAAASA) and MHC class I tetramers TB10.4 (H-2K(b) 4-11, sequence: IMYNYPAM) were obtained from the National Institutes of Health Tetramer Core Facility. Cell sorting was performed on a FACS Aria (BD Biosciences). Sorted cells were collected in complete media, spun down, resuspended in TRIzol, and frozen at −80°C overnight prior to RNA isolation. Samples for flow cytometry were fixed in 2% paraformaldehyde solution in PBS and analyzed using a LSRII flow cytometer (BD Biosciences) and FlowJo software (Tree Star, Inc.).

### Bulk RNA-sequencing and Analysis

RNA isolation was performed using TRIzol (Invitrogen), two sequential chloroform extractions, Glycoblue carrier (Thermo Fisher), isopropanol precipitation, and washes with 75% ethanol. RNA was quantified with the Bioanalyzer RNA 6000 Pico Kit (Agilent). cDNA libraries were constructed using the SMARTer Stranded Total RNA-Seq Kit (v2) – Pico Input Mammalian (Clontech) following the manufacturer’s instructions. Libraries were amplified and then sequenced on an Illumina NextSeq (2 x 76, paired-end (sorted BAL cells) or 2 x 151, paired-end (ex vivo stimulation samples)). Stranded paired-end reads were preprocessed: The first three nucleotides of R2 were removed as described in the SMARTer Stranded Total RNA-Seq Kit – Pico Input Mammalian User Manual (v2: 063017) and read ends consisting of more than 66% of the same nucleotide were removed). The remaining read pairs were aligned to the mouse genome (mm10) + Mtb H37Rv genome using the gsnap aligner (63) (v. 2018-07-04) allowing for novel splicing. Concordantly mapping read pairs (∼20 million / sample) that aligned uniquely were assigned to exons using the 23iocond program and gene definitions from Ensembl Mus_Musculus GRCm38.78 coding and non-coding genes. Genes with low expression were filtered using the “filterByExpr” function in the edgeR package (64). Differential expression was calculated using the “edgeR” package (64) from 23ioconductor.org. False discovery rate was computed with the Benjamini-Hochberg algorithm. Hierarchical clusterings were performed in R using ‘Tsclust’ and ‘hclust’ libraries. Heat map and scatterplot visualizations were generated in R using the ‘heatmap.2’ and ‘ggplot2’ libraries, respectively.

### Gene Set Enrichment Analysis (GSEA)

Input data for GSEA consisted of lists, ranked by −log(p-value), comparing RNAseq expression measures of target samples and naïve controls including directionality of fold-change. Mouse orthologs of human Hallmark genes were defined using a list provided by Molecular Signatures Database (MsigDB) (65). GSEA software was used to calculate enrichment of ranked lists in each of the respective hallmark gene lists, as described previously (66). A nominal p-value for each ES is calculated based on the null distribution of 1,000 random permutations. To correct for multiple hypothesis testing, a normalized enrichment score (NES) is calculated that corrects the ES based on the null distribution. A false-discovery rate (FDR) is calculated for each NES. Leading edge subsets are defined as the genes in a particular gene set that are part of the ranked list at or before the running sum reaches its maximum value.

### Ingenuity Pathway Analysis (IPA)

IPA (QIAGEN) was used to identify enriched pathways for differentially expressed genes between naïve and Mtb-infected AMs (cut-off values: FDR < 0.01, |FC| > 2). The top 20 canonical pathways with enrichment score p-value < 0.05 with greater than 10 gene members are reported.

### Single cell RNA-sequencing

BAL from 10 mice per condition was pooled for each sample, with two independent replicates per condition. Samples were prepared for methanol fixation following protocol “CG000136 Rev. D” from 10X Genomics (67). Briefly, samples were filtered with 70 μm filters and red blood cells were lysed with ACK lysis buffer. Samples were resuspended in 1 mL ice-cold DPBS using a wide-bore tip and transferred to a 1.5 mL low-bind Eppendorf tube. Samples were centrifuged at 700 × g for 5 minutes at 4°C. Supernatant was carefully removed with a p1000 pipette, and the cell pellet was washed two more times with DPBS, counted, and resuspended in 200 μL ice-cold DPBS/1 × 10^6^ cells. 800 μL of ice-cold methanol was added drop-wise for a final concentration of 80% methanol. Samples were incubated at −20°C for 30 minutes and then stored at −80°C for up to 6 weeks prior to rehydration. For rehydration, frozen samples were equilibrated to 4°C, centrifuged at 1,000 × g for 10 minutes at 4°C, and resuspended in 50 μL of Wash-Resuspension Buffer (0.04% BSA + 1mM DTT + 0.2U/μL Protector RNAase Inhibitor in 3× SSC buffer) to achieve ∼1,000 cells/μL (assuming 75% sample loss).

### Single cell RNA-sequencing Analysis

Libraries were prepared using the Next GEM Single Cell 3_ Reagent Kits v3.1 (Dual Index) (10X Genomics) following the manufacturer’s instructions. Raw sequencing data were aligned to the mouse genome (mm10) and UMI counts determined using the Cell Ranger pipeline (10X Genomics). Data processing, integration, and analysis was performed with Seurat v.3 (68). Droplets containing less than 200 detected genes, more than 4000 detected genes (doublet discrimination), or more than 5% mitochondrial were discarded. Genes expressed by less than 3 cells across all samples were removed. Unbiased annotation of clusters using the Immgen database (69) as a reference was performed with “SingleR” package (70). Pseudotime analysis was performed using the “SeuratWrappers” and “Monocle3” R packages (71). Data visualization was performed with the “Seurat”, “tidyverse”, “cowplot”, and “viridis” R packages.

### Alveolar Macrophage *Ex Vivo* Stimulation

AMs were isolated by bronchoalveolar lavage and pooled from 5 mice per group. Cells were plated at a density of 5 x 10^4^ cells/well (96-well plate) in complete RPMI (RPMI plus FBS (10%, VWR), L-glutamine (2mM, Invitrogen), and Penicillin-Streptomycin (100 U/ml; Invitrogen) and allowed to adhere overnight in a 37°C humidified incubator (5% CO_2_). Media with antibiotics and non-adherent cells were washed out prior to stimulation. AM were stimulated with LPS (LPS from Salmonella Minnesota, List Biologicals, #R595, 10 ng/ml), Pam3Cys (Pam3CSK4, EMC Microcollections, GmbH, 10 ng/ml), or H37Rv (effective MOI ∼2:1). H37Rv was prepared by culturing from frozen stock in 7H9 media at 37°C for 48 hours to O.D. of 0.1-0.3. The final concentration was calculated based on strain titer and bacteria was added to macrophages for two hours. Cultures were then washed three times to remove extracellular bacteria. Cell cultures were washed once in PBS after 6 hours to remove dead cells and collected in TRIzol for RNA isolation via chloroform/isopropanol extraction or collected after 20 hours for flow cytometry and ICS.

### Filtering for IFN dependent and independent gene sets

“IFN dependent” and “IFN independent” gene sets were generated from data from Olson et al (38), using the following filters starting from a total of 1,233 genes up-regulated in H37Rv-stimulated WT BMDM with average CPM >1, log_2_ fold change > 1 and FDR < 0.01:

> *“IFN dependent*” = H37Rv-stimulated IFNAR^−/−^ BMDM: log_2_ fold change < 1 AND H37Rv-stimulated WT vs IFNAR^−/−^: log_2_ fold change > 2 = **339 genes**
>
> *“IFN independent”* = H37Rv-stimulated IFNAR^−/−^ BMDM: log_2_ fold change > 1, FDR < 0.01 AND H37Rv-stimulated WT vs IFNAR^−/−^: log_2_ fold change < 2 = **352 genes**

### qRT-PCR

Quantitative PCR reactions were carried out using TaqMan primer probes (ABI) and TaqMan Fast Universal PCR Master Mix (ThermoFisher) in a CFX384 Touch Real-Time PCR Detection System (BioRad). Data were normalized by the level of EF1a expression in individual samples.

### Statistical Analyses

RNA-sequencing was analyzed using the edgeR package from Bioconductor.org and the false discovery rate was computed using the Benjamini-Hochberg algorithm. All other data are presented as mean ± SEM and analyzed by one-way ANOVA (95% confidence interval) with Tukey post-test (for comparison of multiple conditions). Statistical analysis and graphical representation of data was performed using either GraphPad Prism v6.0 software or R. PCA plots generated using “Prcomp” and “Biplot” packages. Venn diagrams and gene set intersection analysis was performed using Intervene (72). p-values, * p < 0.05, ** p < 0.01, *** p < 0.001.

## Supporting information

Summary figure

## Acknowledgements

We thank the Animal Care staff at Seattle Children’s Research Institute and University of Massachusetts Amherst, Pamela Troisch and the Next Gen Sequencing core at the Institute for Systems Biology. The authors acknowledge Research Scientific Computing at Seattle Children’s Research Institute for providing HPC resources that have contributed to the research results reported within this paper. We thank members of the Aderem, Urdahl, and Rothchild labs for helpful discussions.

## Author contributions

D.M., A.C.R, J.N., A.H.D., K.U., and A.A. designed the experiments. A.C.R., D.M., A.J., T.M., P.L., M.C., L.P. conducted the experiments. A.H.D., A.C.R., M.M. performed computational analyses. A.C.R., A.H.D. wrote the paper.

## Data and materials availability

Raw and processed RNA-sequencing data can be accessed from the National Center for Biotechnology Information (NCBI) Gene Expression Omnibus (GEO) database under accession number GSE212205 [https://www.ncbi.nlm.nih.gov/geo/query/acc.cgi?acc=GSE212205]. (*Submission currently private)*

---

**Figure S1 (related to Figure 1): Flow cytometry gating schemes.** Gating strategies for myeloid (A) and T cell (B) analysis.

**Figure S2 (related to Figure 1): *Mycobacterium* exposure provides protection against standard low-dose H37Rv aerosol challenge.** A) Lung, spleen, and lung-draining lymph node (LN) CFU in control mice at deposition, day 10, 12, 14, and 28. B-E) Summary plots of ΔCFU (log) in lung, spleen, and LN following low-dose infection with H37Rv at day 10 (B), day 12 (C), day 14 (D), and day 28 (E). *p < 0.05, **p < 0.01, ***p < 0.001. One-way ANOVA with Tukey post-test. Data compiled from 2-3 independent experiments per condition, with 5 mice per group for each experiment.

**Figure S3 (related to Figure 2): Top 20 Canonical Pathways by Ingenuity Pathway Analysis for up-regulated genes by Mtb-infected alveolar macrophages**. IPA analysis for Mtb-infected AMs from control, scBCG, and coMtb mice 24 hours following high dose mEmerald-H37Rv infection. Data representative of 3 independent experiments per condition.

**Figure S4 (related to Figure 2): Transcriptional changes to naive alveolar macrophages following *mycobacterium* exposure by bulk RNA-sequencing.** A-B) Volcano plots depicting changes in baseline gene expression of naive AMs from scBCG (A) and coMtb(B) mice compared to naive AMs from control mice. Significantly changed genes (FDR < 0.05, |FC| > 2) highlighted and labeled. C) Gene expression for innate receptors and adaptors of interest, log_2_ fold change, unstimulated AMs from scBCG and coMtb mice compared to unstimulated AMs from control mice. * FDR < 0.01. Compiled from 2 independent experiments for each condition.

**Figure S5 (related to Figure 3): Flow analysis of BAL samples prepared for 10X single-cell RNA-sequencing.** Percentage of each population (AM, lymphocytes, eosinophils, MDM, other CD45^+^) out of CD45^+^ ZV^−^. AM = Siglec F^+^ CD64^+^, Eosinophils = Siglec F^+^ CD64^−^, lymphocytes = CD3/CD19^+^, MDM = Siglec F^−^ CD64^+^, other CD45^+^ = CD3^−^ CD19^−^ Siglec F^−^ CD64^−^. Note: One of the two coMtb samples analyzed by flow cytometry did not have an accompanying 10X sample. The second coMtb 10X sample was processed separately without flow analysis.

**Figure S6 (related to Figure 3): Top 10 genes differentially expressed for each of 11 macrophage subclusters** Heatmap of genes that are most differentially expressed for each of 11 clusters with all other clusters. Genes filtered with log fold change threshold of > 0.25 and minimum percentage expression of 25% of cells. All genes but one (*Gsto1*) had an adjusted p-value of < 1.0×10-5 . *Five genes (*Fabp4, Fabp5, Stmn1, Mki67, Cbr2*) met this criterion for more than one cluster, grouped with the more abundant cluster. Data is compiled from two independent experiments, 3 conditions each, for a total of 6 samples.

**Figure S7 (related to Figure 3) UMAP gene expression plots for genes associated with macrophage subcluster 3 and found in AM_2 (Pisu et al).** Genes associated with mitochondrial oxidative phosphorylation (*mt-Co1, mt-Cytb, mt-Nd2*), chromatin remodeling (*Ankrd11, Baz1a*), macrophage-associated transcription factors (*Cebpb, Zeb2, Bhlhe40, Hif1a*), and CARD9 signaling (*Malt1, Bcl10*). Data is compiled from two independent experiments with 3 conditions each, for a total of 6 samples.

**Figure S8 (related to Figure 4): UMAP gene expression plots of cluster and lineage marker genes of interest for T cell subclusters.** Data is compiled from two independent experiments with 3 conditions each for a total of 6 samples.

**Figure S9 (related to Figure 5): Gene expression of alveolar macrophages from *ex vivo* stimulations.** A) AMs were stimulated for 6 hours with Pam3Cys (10 ng/ml), LPS (10 ng/ml), or H37Rv (effective MOI ∼2:1). Volcano plots depict fold change (log_2_) and P-value (-log_10_) for each stimulation condition for each of the three groups (control scBCG, coMtb) compared to the respective unstimulated controls. DEG (p-value < 0.001; |fold change| > 2) highlighted and labeled, space permitting. B) Baseline gene expression for innate receptors and adaptors of interest, log_2_ fold change, unstimulated AMs from scBCG and coMtb mice compared to unstimulated AMs from control mice. Compiled from 3 independent experiments.

**Figure S10 (related to Figure 5): Cell-intrinsic changes in alveolar macrophage response is retained 23 weeks following vaccination.** Gene expression of *Mx1, Cxcl10, Il1b, Cxcl2, Irf7,* and *Il6* as measured by qPCR in AMs isolated by BAL from mice 8 and 23 weeks following scBCG vaccination and from age-matched controls, with and without LPS (10 ng/ml) stimulation. Data is representative of technical AM duplicates from a single experiment.

Table S1: RNA-Sequencing data for alveolar macrophages 24 hours following high dose H37Rv-mEmerald challenge from scBCG mice

Table S2: Top differentially expressed genes for individual clusters for macrophage, T cell, and dendritic cell sub-cluster analysis

Table S3: RNA-Sequencing data for *ex vivo* stimulated alveolar macrophages

Table S4: IFN-independent and IFN-dependent genes based on WT and IFNAR^−/−^ BMDM RNA-seq data

