## Supplemental Figures 1-10 for "Exposure to *mycobacterium* remodels alveolar macrophages and the early innate response to *Mycobacterium tuberculosis* infection"

**Supplementary Materials**

Fig. S1: Lung flow cytometry gating schemes

Fig. S2: *Mycobacterium* exposure provides protection against standard low-dose H37Rv aerosol challenge

Fig. S3: Top 20 Canonical Pathways by Ingenuity Pathway Analysis for up-regulated genes by Mtb-infected alveolar macrophages

Fig. S4: Transcriptional changes to naïve alveolar macrophages following *mycobacterium* exposure by bulk RNA-sequencing

Fig. S5: Flow analysis of BAL samples prepared for 10X single-cell RNA-sequencing

Fig. S6: Top 10 genes differentially expressed for each of 11 macrophage subclusters

Fig. S7: UMAP gene expression plots for genes associated with macrophage subcluster 3 and found in AM_2 (Pisu et al)

Fig. S8: UMAP gene expression plots of cluster and lineage marker genes of interest for T cell subclusters

Fig. S9: Gene expression of alveolar macrophages from *ex vivo* stimulations

Fig. S10: Cell-intrinsic changes in alveolar macrophage response is retained 23 weeks following vaccination

Table S1: RNA-Sequencing data for alveolar macrophages 24 hours following high dose H37Rv-mEmerald challenge from scBCG mice

Table S2: Top differentially expressed genes for individual clusters for macrophage, T cell, and dendritic cell sub-cluster analysis

Table S3: RNA-Sequencing data for ex vivo stimulated alveolar macrophages

Table S4: IFN-independent and IFN-dependent genes based on WT and IFNAR$^{-/-}$ BMDM RNA-seq data

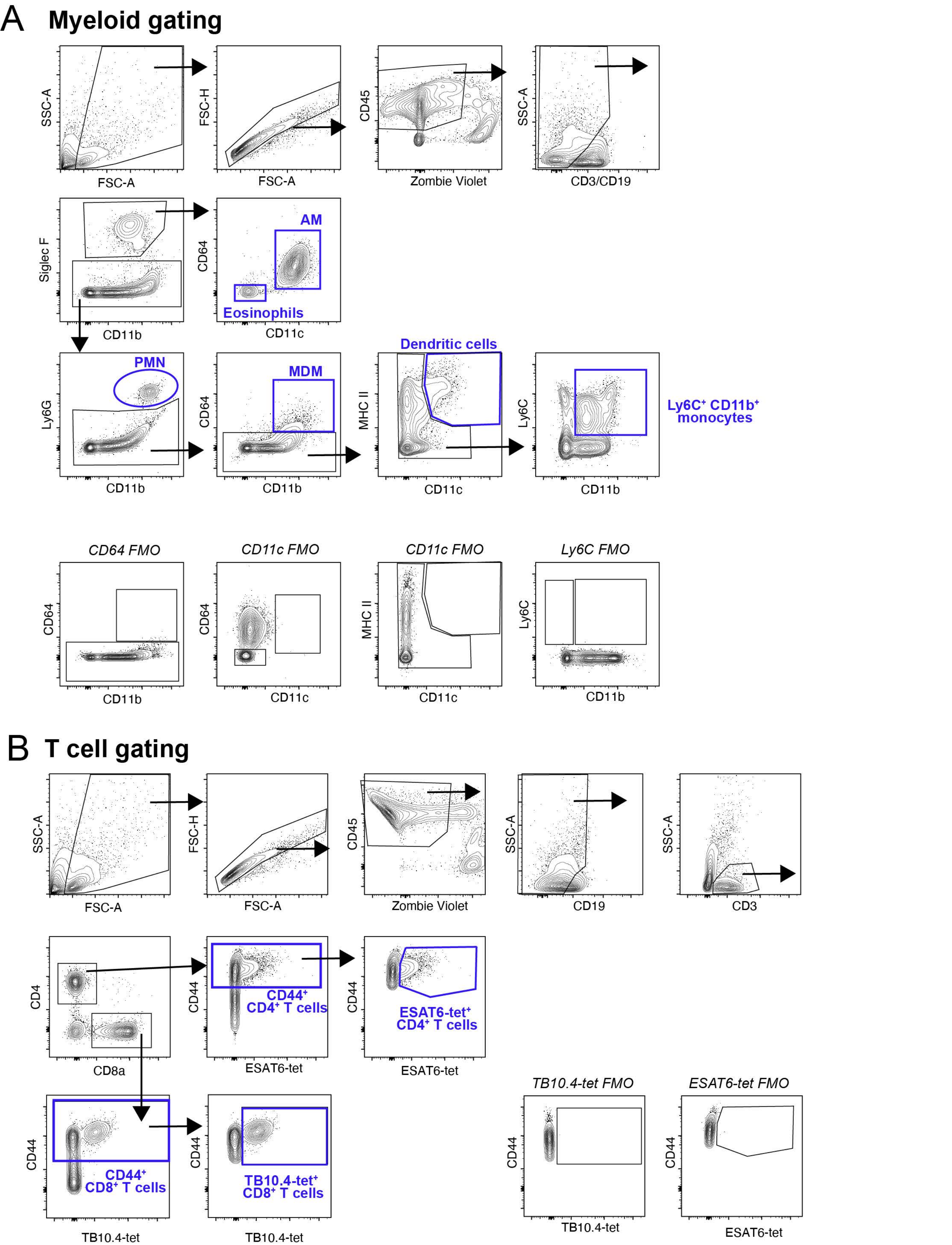


**Figure S1 (related to Figure 1). Flow cytometry gating schemes**. Gating strategies for myeloid (A) and T cell (B) analysis.

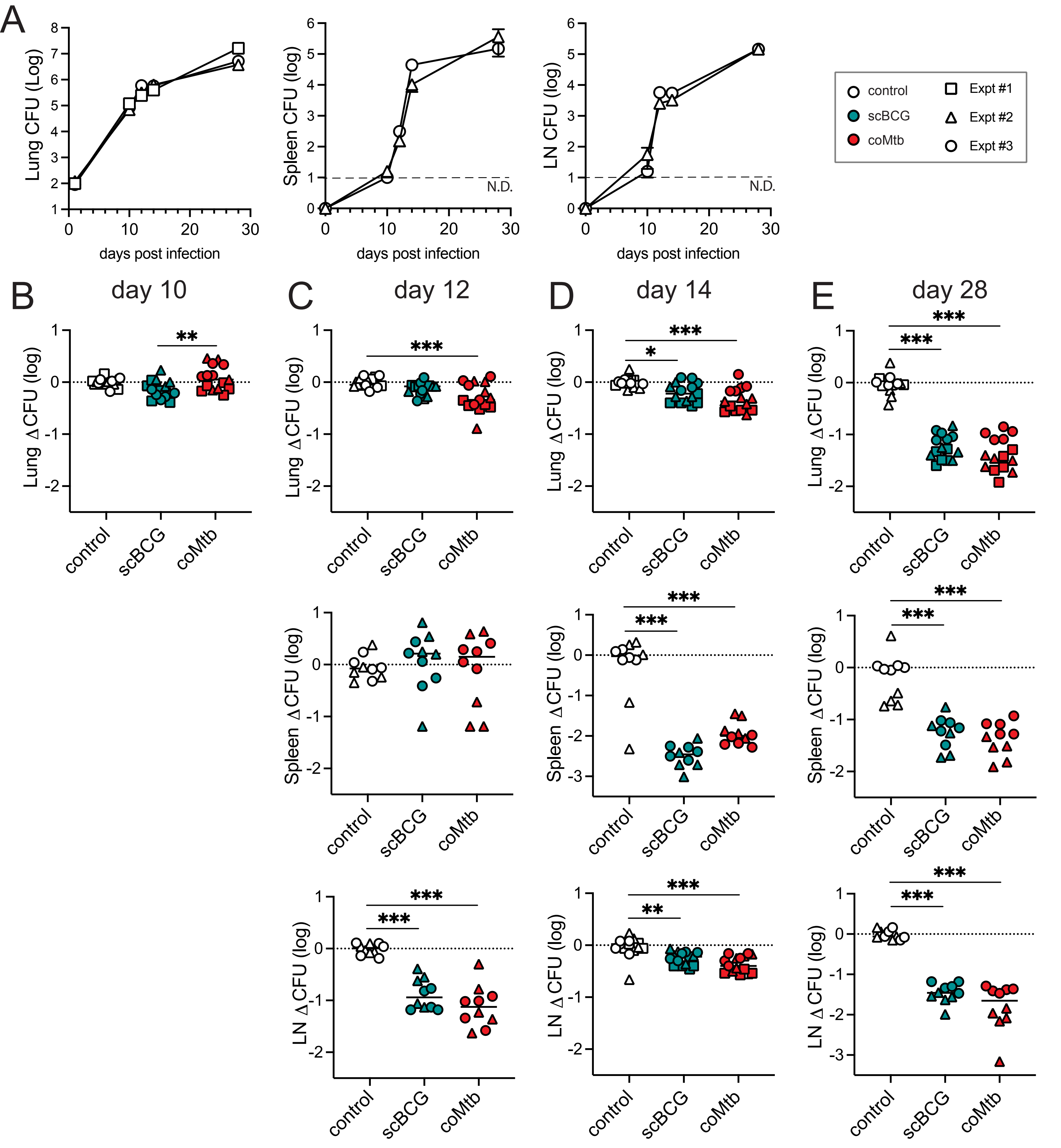


**Figure S2 (related to Figure 1): *Mycobacterium* exposure provides protection against standard low-dose H37Rv aerosol challenge.** A) Lung, spleen, and lung-draining lymph node (LN) CFU in control mice at deposition, day 10, 12, 14, and 28. B-E) Summary plots of ΔCFU (log) in lung, spleen, and LN following low-dose infection with H37Rv at day 10 (B), day 12 (C), day 14 (D), and day 28 (E). $^{*}p < 0.05$, $^{**}p < 0.01$, $^{***}p < 0.001$. One-way ANOVA with Tukey post-test. Data compiled from 2-3 independent experiments per condition, with 5 mice per group for each experiment.

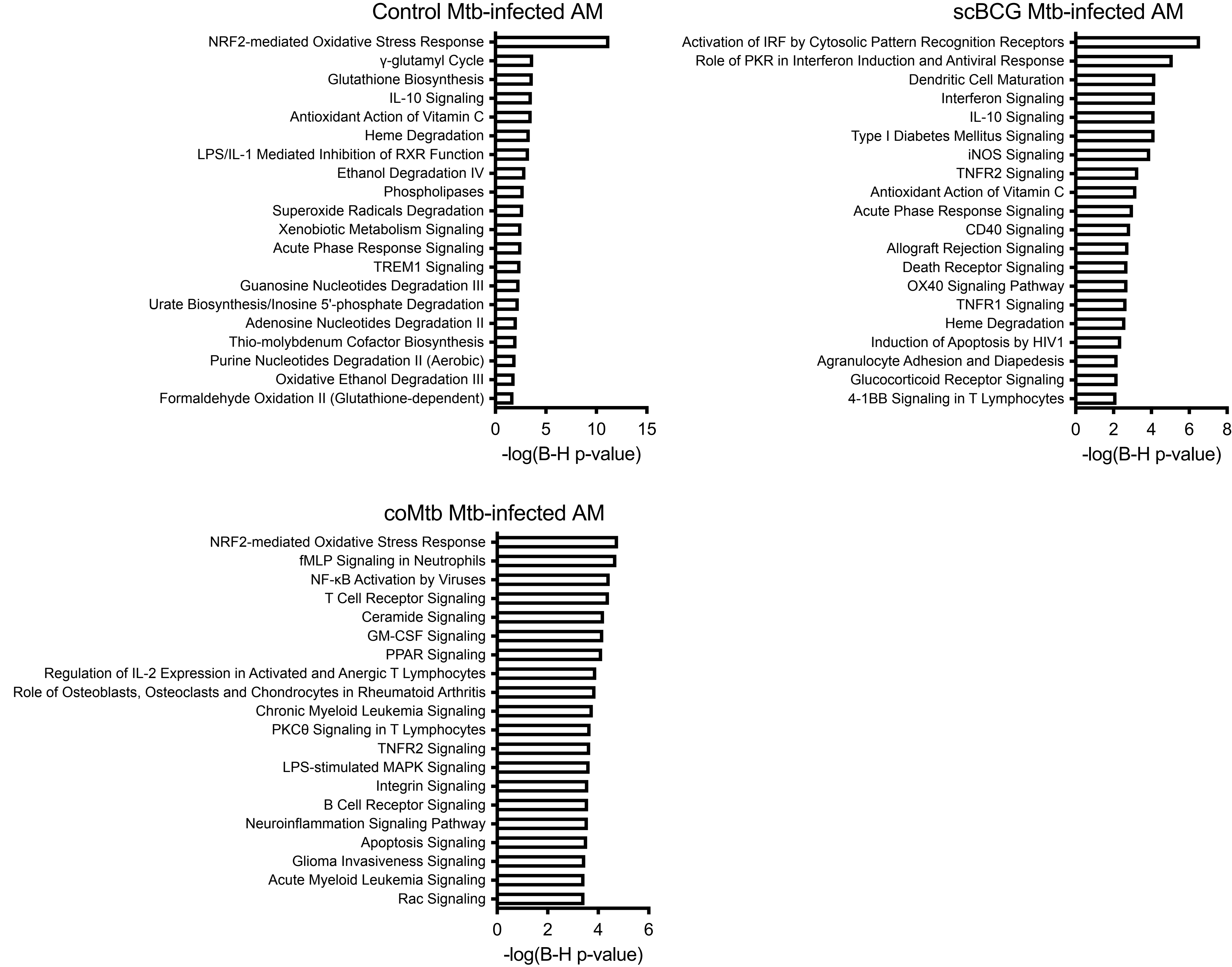


**Figure S3 (related to Figure 2): Top 20 Canonical Pathways by Ingenuity Pathway Analysis for up-regulated genes by Mtb-infected alveolar macrophages.** IPA analysis for Mtb-infected AMs from control, scBCG, and coMtb mice 24 hours following high dose mEmerald-H37Rv infection. Data representative of 3 independent experiments per condition.

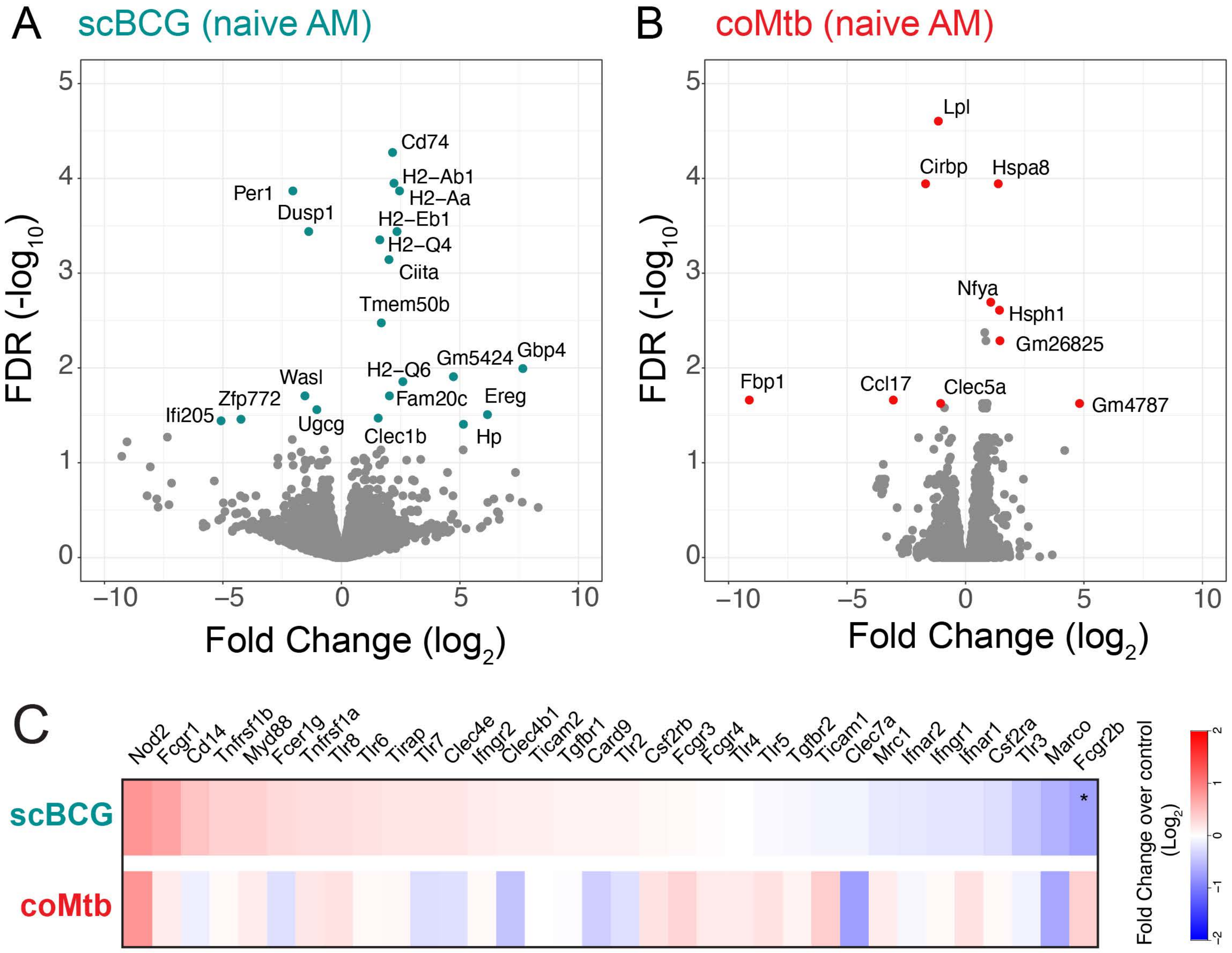


**Figure S4 (related to Figure 2): Transcriptional changes to naive alveolar macrophages following *myco-bacterium* exposure by bulk RNA-sequencing.** A-B) Volcano plots depicting changes in baseline gene expression of naive AMs from scBCG (A) and coMtb(B) mice compared to naive AMs from control mice. Significantly changed genes (FDR < 0.05, |FC| > 2) highlighted and labeled. C) Gene expression for innate receptors and adaptors of interest, $log_2$ fold change, unstimulated AMs from scBCG and coMtb mice compared to unstimulated AMs from control mice. * FDR < 0.01. Compiled from 2 independent experiments for each condition.

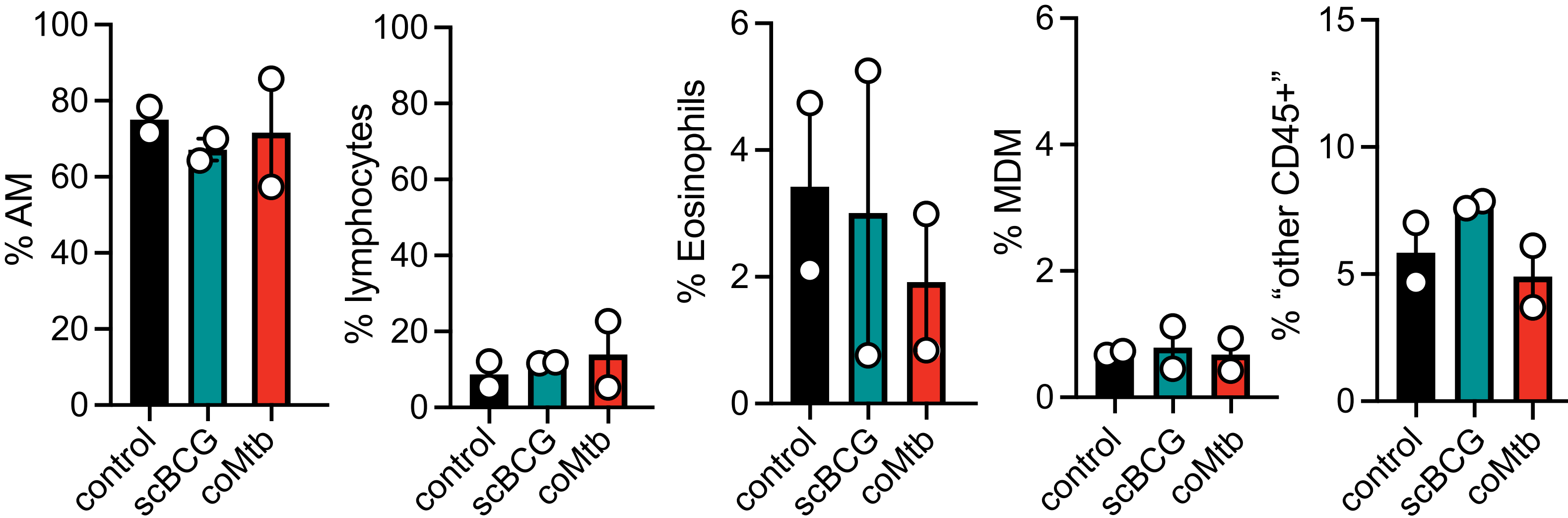


**Figure S5 (related to Figure 3): Flow analysis of BAL samples prepared for 10X single-cell RNA-sequencing.** Percentage of each population (AM, lymphocytes, eosinophils, MDM, other CD45+) out of $CD45^+$ $ZV^-$. AM = Siglec $F^+$ $CD64^+$, Eosinophils = Siglec $F^+$ $CD64^-$, lymphocytes = $CD3/CD19^+$, MDM = Siglec $F^-$ $CD64^+$, other CD45+ = $CD3^-$ $CD19^-$ Siglec $F^-$ $CD64^-$. Note: One of the two coMtb samples analyzed by flow cytometry did not have an accompanying 10X sample. The second coMtb10X sample was processed separately without flow analysis.

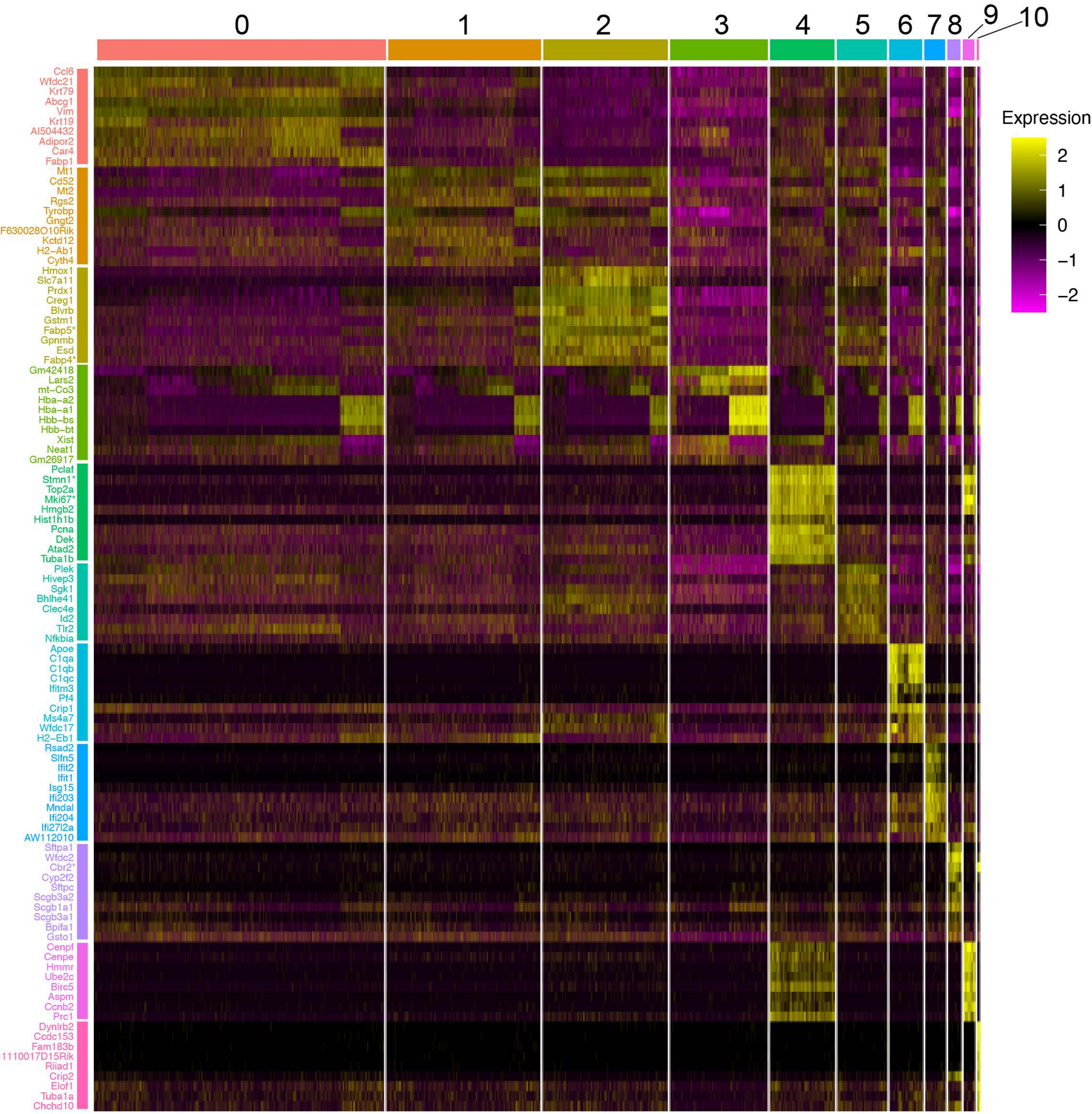


**Figure S6 (related to Figure 3). Top 10 genes differentially expressed for each of 11 macrophage subclusters.** Heatmap of genes that are most differentially expressed for each of 11 clusters with all other clusters. Genes filtered with log fold change threshold of > 0.25 and minimum percentage expression of 25% of cells. All genes but one (*Gsto1*) had an adjusted p-value of < $1.0 \times 10^{-5}$ . *Five genes (*Fabp4, Fabp5, Stmn1, Mki67, Cbr2*) met this criteria for more than one cluster, grouped with the more abundant cluster. Data is compiled from two independent experiments, 3 conditions each, for a total of 6 samples.

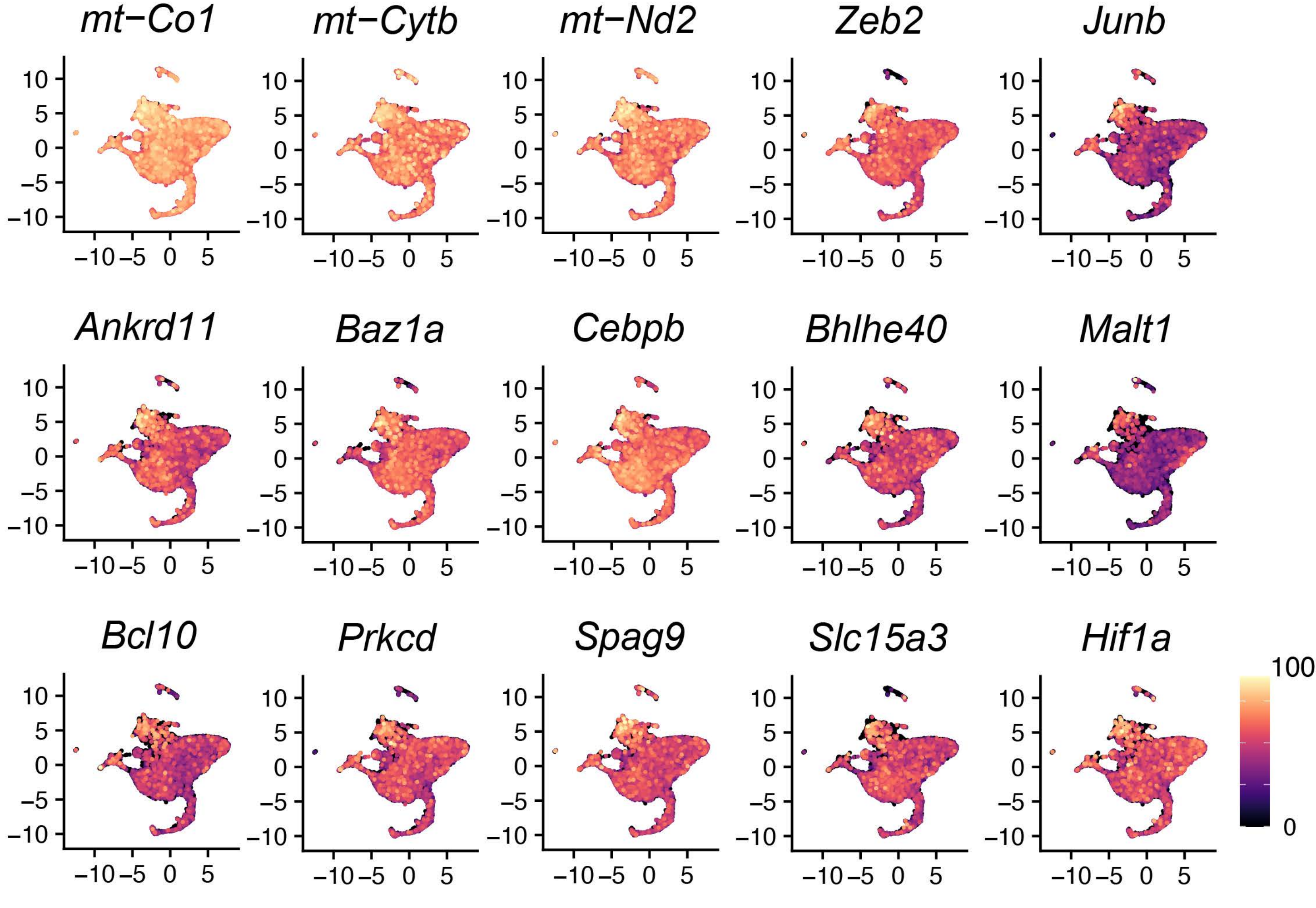


**Figure S7 (related to Figure 3) UMAP gene expression plots for genes associated with macrophage subcluster 3 and found in AM_2 (Pisu et al).** Genes associated with mitochondrial oxidative phosphorylation (*mt-Co1, mt-Cytb, mt-Nd2*), chromatin remodeling (*Ankrd11, Baz1a*), macrophage-associated transcription factors (*Cebpb, Zeb2, Bhlhe40, Hif1a*), and CARD9 signaling (*Malt1, Bcl10)*. Data is compiled from two independent experiments with 3 conditions each, for a total of 6 samples.

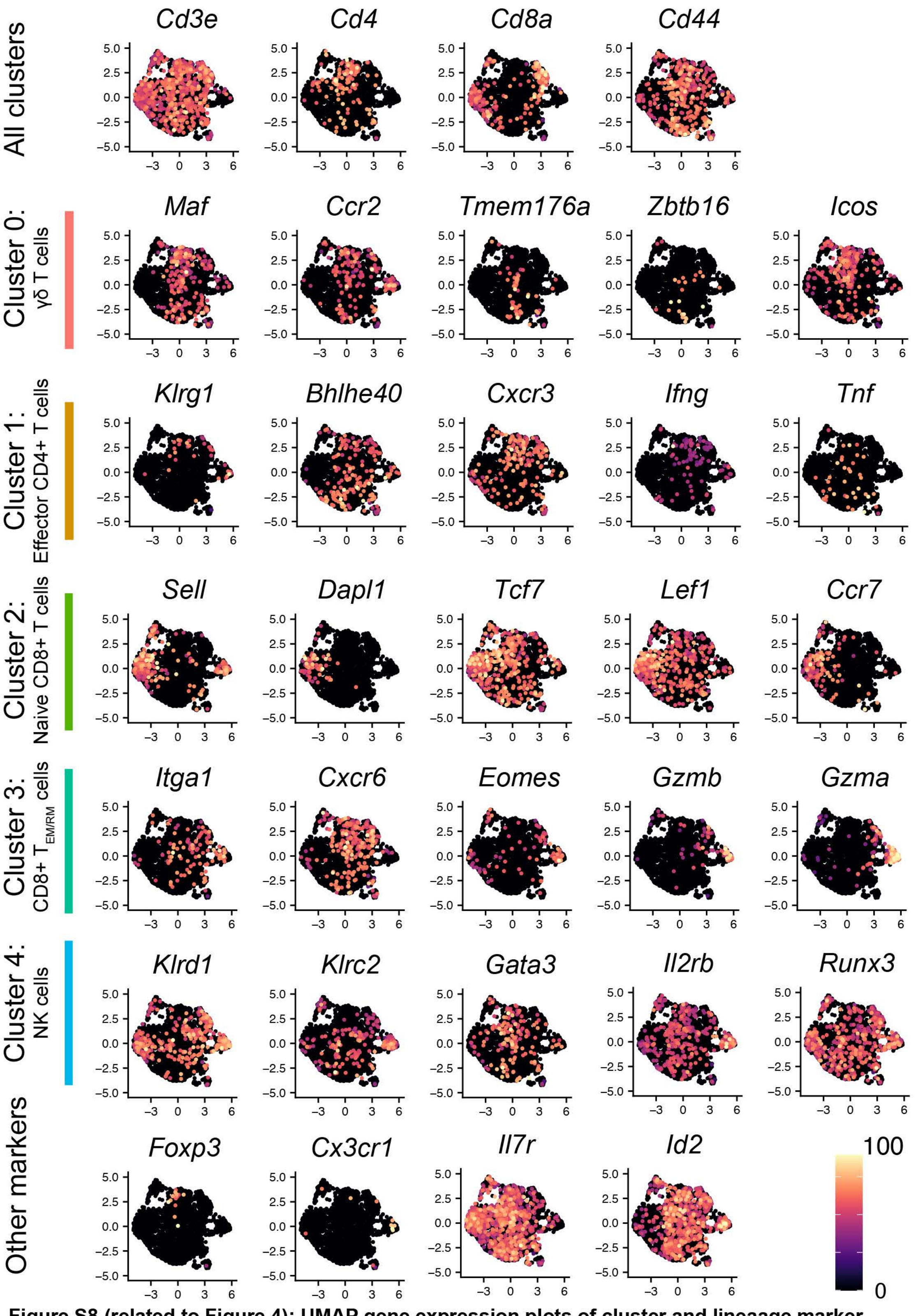


**Figure S8 (related to Figure 4): UMAP gene expression plots of cluster and lineaage marker genes of interest for T cell subclusters.** Data is compiled from 2 independent experiments with 3 conditions each for a total of 6 scRNAseq BAL samples.

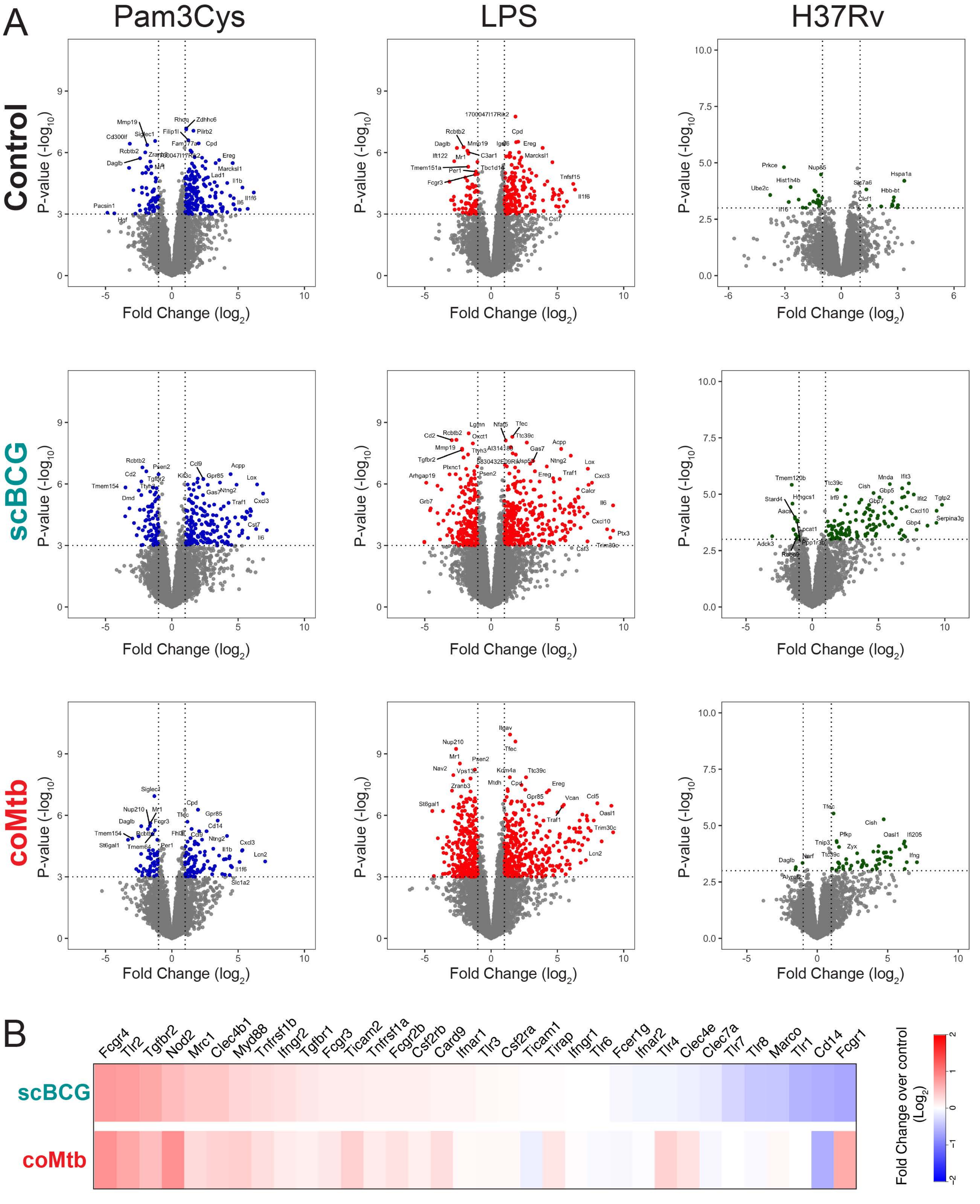


**Figure S9 (related to Figure 5): Gene expression of alveolar macrophages from *ex vivo* stimulations.** A) AMs were stimulated for 6 hours with Pam3Cys (10 ng/ml), LPS (10 ng/ml), or H37Rv (effective MOI ~2:1). Volcano plots depict fold change ($\log_2$) and P-value ($-\log_{10}$) for each stimulation condition for each of the three groups (control scBCG, coMtb) compared to the respective unstimulated control. DEG (p-value < 0.001; |fold change| > 2) highlighted and labeled, space permitting. B) Baseline gene expression for innate receptors and adaptors of interest, $\log_2$ fold change, unstimulated AMs from scBCG and coMtb mice compared to unstimulated AMs from control mice. Compiled from 3 independent experiments.

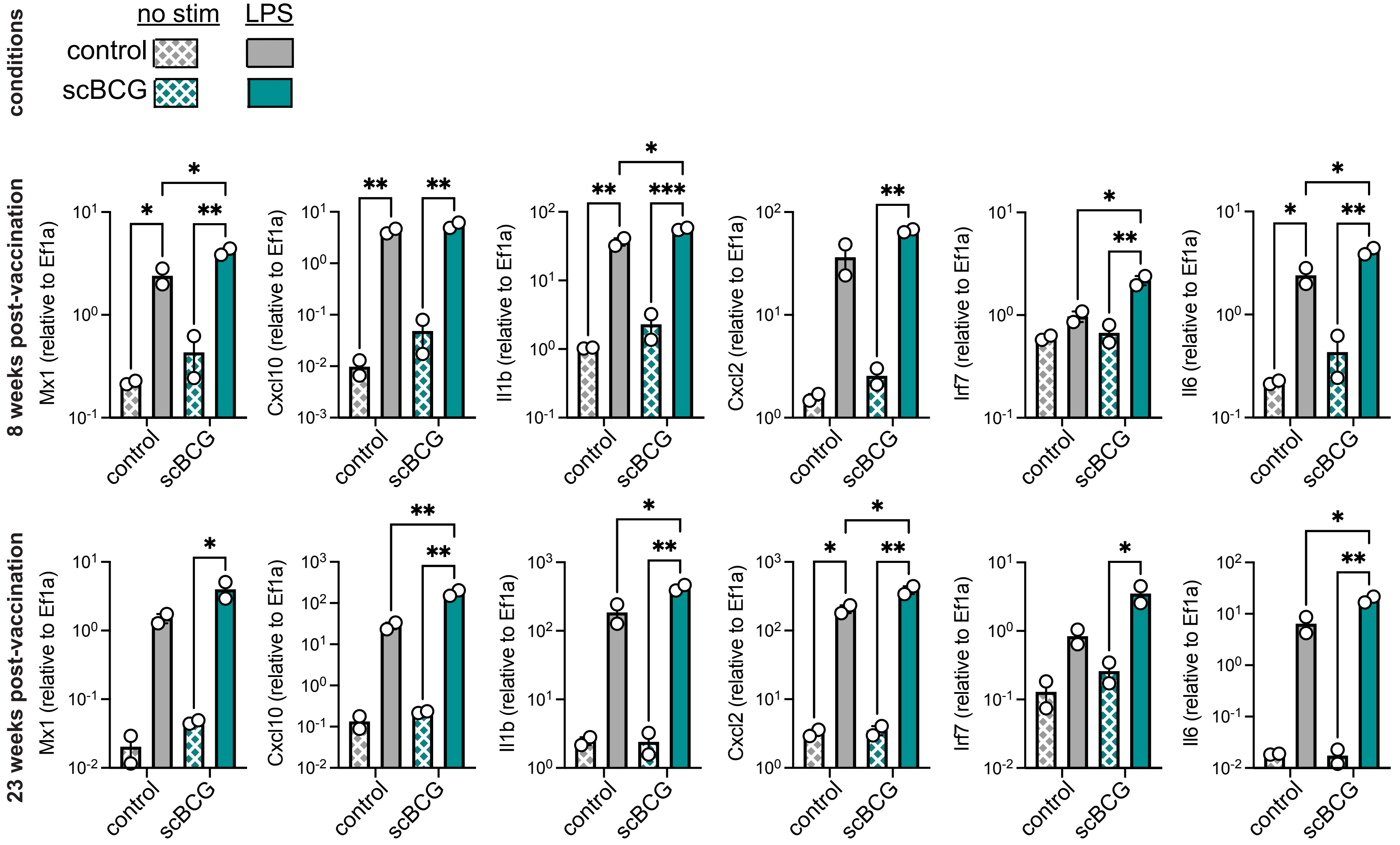


**Figure S10 (related to Figure 5): Cell-intrinsic changes in alveolar macrophage response is retained 23 weeks following vaccination.** Gene expression of *Mx1, Cxcl10, Il1b, Cxcl2, Irf7,* and *Il6* as measured by RT-qPCR in alveolar macrophages isolated by BAL from mice 8 and 23 weeks following scBCG vaccination and from age-matched controls, with and without LPS (10 ng/ml) stimulation. Data is representative of technical AM duplicates from a single experiment.
