## Supplementary figures and images for "Exposure to *mycobacterium* remodels alveolar macrophages and the early innate response to *Mycobacterium tuberculosis* infection"

### Summary figure

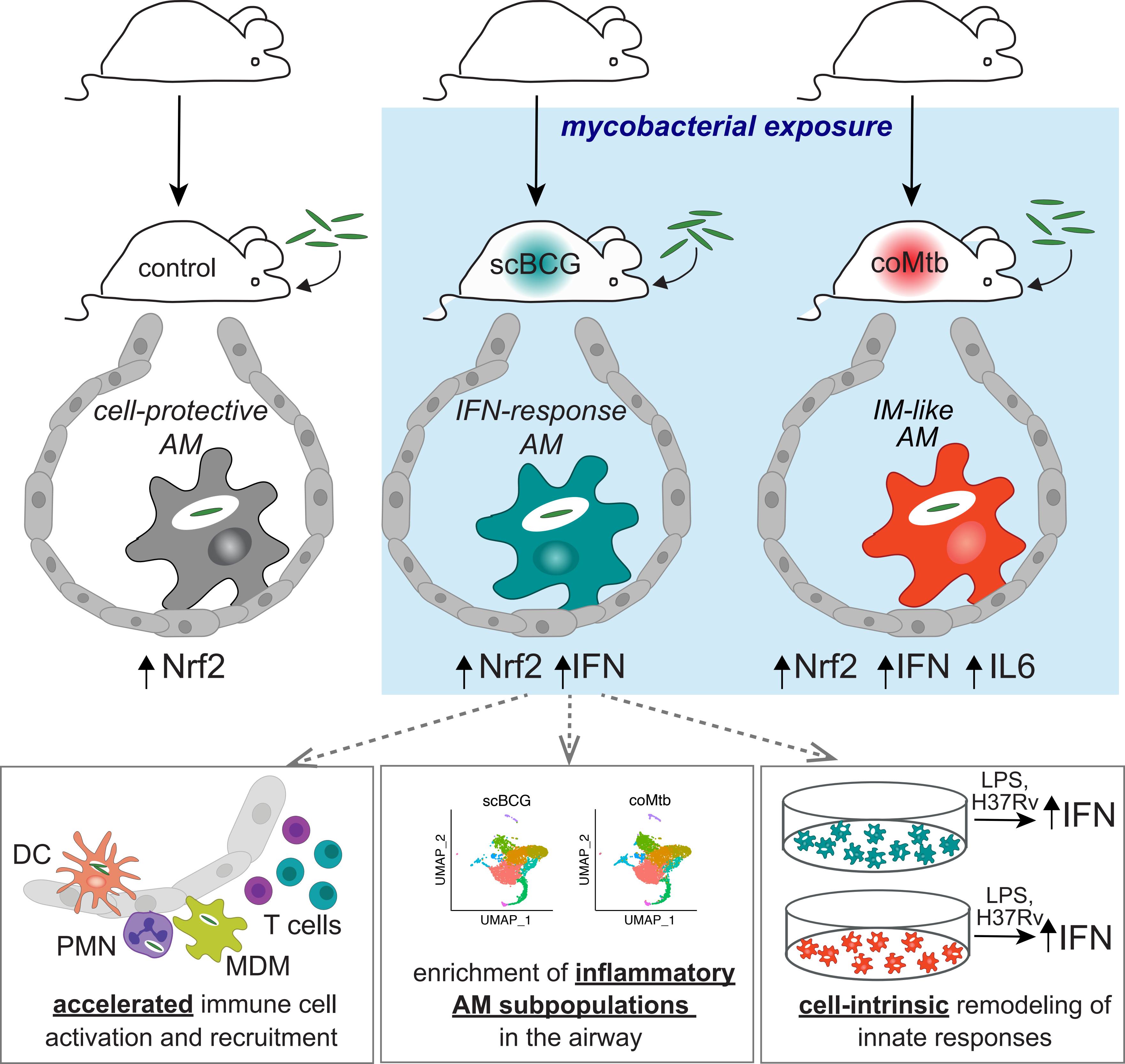
